# Epigenetic therapy remodels the immune synaptic cytoskeleton to potentiate cancer susceptibility to γδ T cells

**DOI:** 10.1101/2020.04.30.069955

**Authors:** Rueyhung R. Weng, Hsuan-Hsuan Lu, Chien-Ting Lin, Chia-Chi Fan, Rong-Shan Lin, Tai-Chung Huang, Shu-Yung Lin, Yi-Jhen Huang, Yi-Hsiu Juan, Yi-Chieh Wu, Zheng-Ci Hung, Chi Liu, Xuan-Hui Lin, Wan-Chen Hsieh, Tzu-Yuan Chiu, Jung-Chi Liao, Yen-Ling Chiu, Shih-Yu Chen, Chong-Jen Yu, Hsing-Chen Tsai

## Abstract

γδ T cells are a distinct subgroup of T cells that bridge the innate and adaptive immune systems and can attack cancer or virus-infected cells in an MHC-unrestricted manner. Despite its antitumor ability in both autologous and allogeneic settings, earlier trials of adoptive γδ T cell transfer in solid tumors had limited success due to limitations in cell expansion and the lack of a strategy to modulate tumor lytic interactions between γδ T and cancer cells. Here, we show through quantitative surface proteomics and gene enrichment analyses that DNA methyltransferase inhibitors (DNMTis) upregulate multiple surface molecules related to γδ T cell activation in cancer cells. DNMTi treatment of human lung cancer potentiates tumor lysis by *ex vivo*-expanded γδ T cells using a clinical-grade expansion protocol developed by our team to enrich for the Vδ1 subset while preserving their antitumor effector functions. Mechanistically, DNMTis enhance immune synapse formation and stabilize the synaptic cleft to facilitate γδ T-mediated tumor lysis. Through integrated analysis of RNA-seq, DNA methylation, and ATAC-seq, we demonstrate that depletion of DNMTs induces coordinated pattern alterations of immune synaptic-cytoskeletal networks at the cancer side of the immune synapse. In addition, single-cell mass cytometry reveals enrichment of polyfunctional γδ T subsets by DNMTis. Combined DNMTi and adoptive γδ T transfer in a mouse lung cancer model offers a significant survival benefit. Consistently, the DNMTi-associated cytoskeleton signature identifies a subset of lung cancer patients with improved survival. Our results demonstrate that epigenetic mechanisms are crucial for cytoskeletal remodeling in cancer to potentiate immune attack and support a combinatorial strategy of DNMTis and γδ T cell-based immunotherapy in lung cancer management.

**One Sentence Summary:** DNA methyltransferase inhibitors potentiate the killing of lung cancer by γδ T cells through remodeling cytoskeletal-immune synaptic networks.

## INTRODUCTION

DNA methyltransferase inhibitors (DNMTis) are used clinically in treating myelodysplastic syndrome and hematological malignancies. Two FDA-approved DNMTis, decitabine (Dacogen®, DAC) and azacytidine (Vidaza®, AZA), are cytidine analogs that incorporate into DNA/RNA and deplete DNMTs through irreversible binding of the enzymes followed by proteasome degradation (*1*). The loss of DNMTs leads to substantial global gene upregulation by altering epigenetic regulatory processes, including demethylation of DNA or breakdown of repressive complexes that contain DNMTs as scaffold proteins. We previously demonstrated that DNMTis at noncytotoxic doses may produce a memory type of cellular response associated with alterations in several major signaling networks, including immune-related pathways, and exert long-lasting antitumor effects in multiple cancer types (*2*).

Compelling evidence has indicated the immunomodulatory effects of DNMTis on cancer and immune cells, targeting the major histocompatibility complex (MHC) and T cell receptor (TCR) axis (*3, 4*). Studies have shown that DNMTis may augment antitumor immune responses through reexpression of cancer-testis antigens, antigen processing, and antigen presentation machinery, as well as immune checkpoint molecules (*5–7*). DNMTis have also been linked to activation of endogenous retroviral sequences and induction of antiviral interferon responses in cancer cells (*4, 8*), as well as reversing the exhausted phenotype of tumor-infiltrating T cells (*9, 10*). Likewise, knockdown of DNMT1 below a critical threshold may reactivate multiple viral defense- and immune-related genes in colorectal cancers (*11*). All of these findings provide solid molecular rationales for combinatorial therapy of DNMTis and checkpoint inhibitors (*12*). Nevertheless, for patients who are inherently unresponsive to checkpoint blockade or have acquired defects along the TCR-MHC axis following therapy, the added therapeutic benefits of epigenetic therapy may be modest. As various types of cancer immunotherapies targeting surface molecules to exert MHC-unrestricted antitumor immunity, such as γδ T, natural killer (NK), and chimeric antigen receptor (CAR) T cells, are under active development, understanding whether and how DNMTis may reshape surface proteins of cancer cells and augment these immunotherapeutic strategies in patients unsuitable for checkpoint inhibitors is a promising approach.

Here, we show that DNMTis markedly alter the surface proteome of lung cancer cells using stable isotope labeling by amino acids in cell culture (SILAC)-based quantitative proteomics. The data reveal sustained upregulation of multiple immune-related molecules following transient DNMTi treatment at low doses. Many of them are bona fide immune synapse proteins. Gene ontology analysis indicates a high association of γδ T cell activation via the DNMTi-induced surface proteome. Functional studies were conducted to demonstrate that DNMTi treatment of cancer cells significantly enhances immune synapse formation between *ex vivo* expanded allogeneic γδ T cells and cancer cells, potentiating antitumor immunity by γδ T cells. A combined genome-wide analysis of the DNA methylome, mRNA-seq, and Omni-ATAC-seq reveals coordinated epigenetic regulatory patterns of immune synaptic cytoskeleton networks. Our findings support the broad applicability of DNMTis in priming cancer cells for MHC-unrestricted γδ T cell-based therapy.

## RESULTS

### Decitabine upregulates surface immune molecules related to γδ T cell activation

DNMTis have been shown to modulate pathways related to antigen processing/presentation, human leukocyte antigens (HLA), and interferon responses across different cancer types at the transcriptomic level (*6, 7*). Nevertheless, these transcriptomic changes may not fully reflect the alterations of surface proteins that account for susceptibility to immunotherapy. Thus, we sought to obtain a comprehensive profile of surface proteins altered by decitabine (DAC) through isolating cell surface proteins from A549 human lung cancer cells before and after DAC treatment using the EZ-link Sulfo-NHS-SS-biotin-assisted biotinylation method, followed by a SILAC-based quantitative proteomics approach (**Fig. 1A**) (*13*). We employed a low-dose treatment protocol established in our previous study to manifest the drug’s epigenetic effects over cytotoxicity (**Fig. 1B**) (*2*) and identified 666 and 831 Gene Ontology (GO)-annotated surface proteins (corresponding to 8,791 and 11,898 unique peptides) in A549 cells upon 100 nM DAC treatment for 72 hours (D3) and growth in drug-free medium for another 3 days (D3R3), respectively (**Fig. 1C and fig. S1A**). The continued increase in identified surface proteins after drug withdrawal is consistent with our prior report of epigenetic memory effects following transient drug exposure (*2*). Among all the identified proteins, 314 proteins showed sustained upregulation upon DAC treatment by at least 1.4-fold for both D3 and D3R3 (**Fig. 1D**). Many of these are immune-related proteins. In addition to MHC molecules and certain immune checkpoint proteins that were previously known to be upregulated by DAC (*6, 7*), we uncovered a plethora of proteins that participate in innate immunity or MHC-unrestricted immunity, such as MHC class I polypeptide-related sequences A and B (i.e., MICA, MICB), which are ligands of NKG2D on γδ T and NK cells (**Fig. 1E**) (*14*). DAC also upregulates UL16-binding proteins 1, 2, and 3 (i.e., ULBP1, ULBP2, and ULBP3), another group of NKG2D ligands that are expressed in many cancers and in stressed/damaged tissues (**Fig. 1E**) (*15, 16*). These molecules have been identified as targets for tumor immunosurveillance by the innate immune system and may elicit antitumor immunity without the requirement for conventional MHC-restricted antigen presentation (*17*). In addition, the two death receptors (i.e., TRAIL-R1 and TRAIL-R2) for TNF-related apoptosis-inducing ligand (TRAIL) that induce cancer apoptosis as part of immune surveillance (*18*) were significantly upregulated in D3R3 (**Fig. 1E**).

**Fig. 1.**
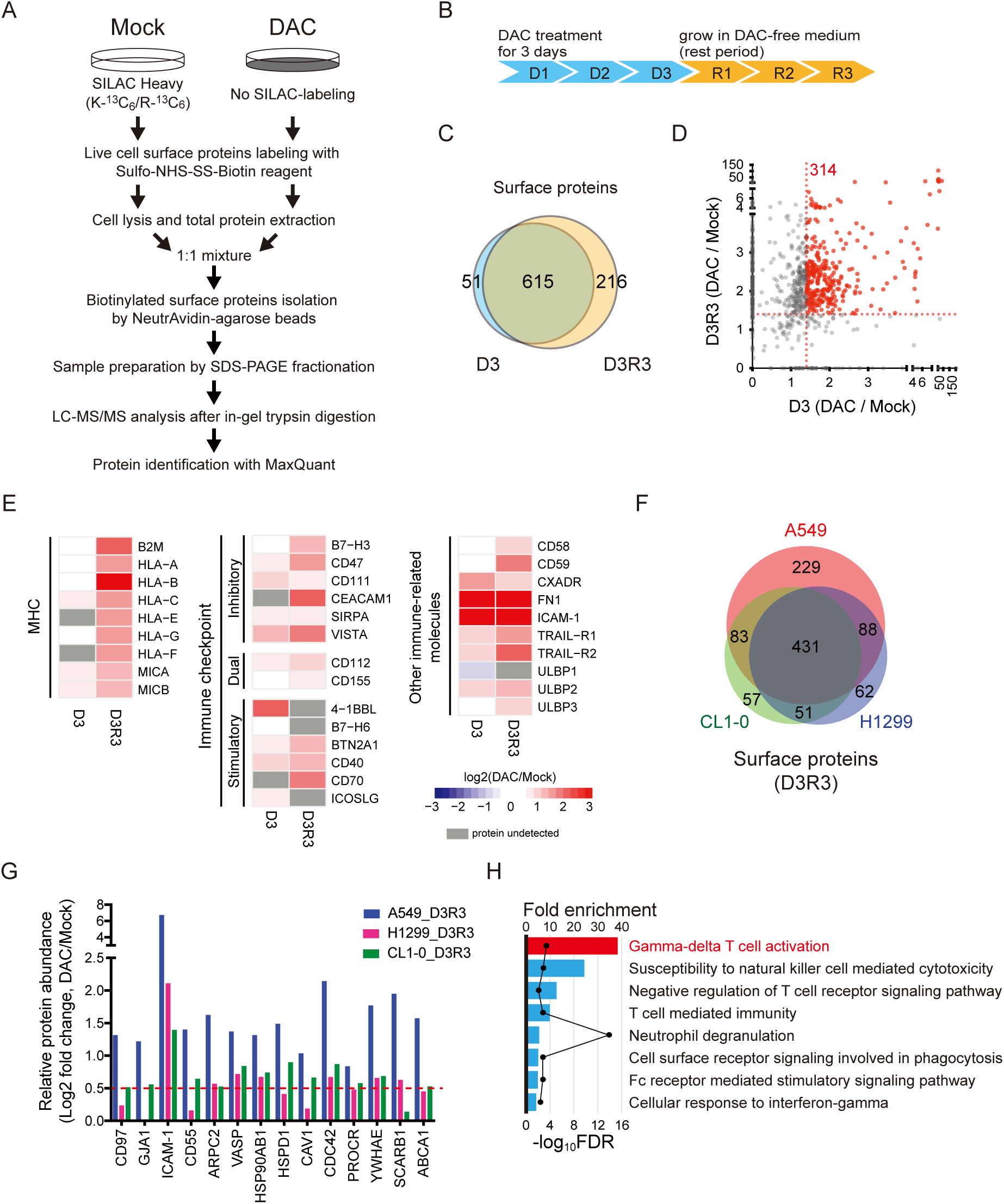
Decitabine (DAC) upregulates surface immune molecules related to γδ T activation. (**A**) Experimental diagram of stable isotope labeling with amino acids in cell culture (SILAC)-based quantitative proteomics on biotinylated surface proteins in mock-treated vs. DAC-treated lung cancer cells. (**B**) Treatment schedule of DAC at 100 nM daily for 72 hours (D3), followed by drug withdrawal for three days (D3R3). (**C**) Venn diagram showing the numbers of surface proteins identified at D3 and D3R3 in A549 cells. (**D**) A scatter plot of proteins upregulated at D3 and D3R3 in A549 cells following decitabine treatment. (**E**) Heatmap showing log2 fold changes of immune-related surface molecules in DAC-treated vs. mock-treated A549 cells at D3 and D3R3. (**F**) Venn diagram showing numbers of surface proteins commonly identified at D3R3 in A549, H1299, and CL1-0 cells. (**G**) Bar graphs showing relative protein abundance of selected surface proteins related to innate immunity in surface proteomes of A549, H1299, and CL1-0 cells following decitabine treatment at D3R3 as compare with mock-treated cells. (**H**) PANTHER gene list analysis on immune-related pathways for proteins upregulated by decitabine at D3R3.

Interestingly, analysis of SILAC-based surfaceomes in another two human lung cancer cell lines, H1299 and CL1-0, revealed highly similar profiles of DAC-induced surface proteins. There were 431 proteins commonly upregulated by DAC in all three lung cancer cell lines for D3R3 (**Fig. 1F and fig. S1B**). The involvement of DAC-induced surface molecules in the innate immune response is also evident, and we mapped these proteins against the innate immune interactomes from InnateDB, an extensively curated database of innate immune pathways and interactions (*19*). We uncovered DAC-mediated innate immune molecules that participate in adhesion/cell-cell interactions (e.g., CD97 and ICAM-1), the cytoskeleton (e.g., ARPC2 and VASP), heat shock protein responses, integrin-associated pathways, and various signal transduction networks (**Fig. 1G**). Next, we performed Gene Ontology (GO) term enrichment using PANTHER Gene List Analysis tools (*20*) on the DAC-induced surface proteome to search for relevant immune pathways. Remarkably, γδ T cell activation was the top enriched pathway among all immune-related processes, followed by natural killer (NK) cell-mediated immunity — two of the key immune cell types involved in innate immune responses against cancer (**Fig. 1H**) (*21, 22*).

### *Ex vivo* expanded human Vδ1-enriched γδ T cells retain antitumor effector functions

Based on the DAC-mediated surfaceome data, we reason that epigenetic therapy has the potential to enhance tumor attack by γδ T cells, a distinct subset of T cells that do not require classical MHC molecules for antigen recognition, and thus is a promising cell type for autologous or allogeneic adoptive cell transfer therapy against various types of cancer in clinical trials (*23*). Two major subsets of human γδ T cells, Vδ1+ (intraepithelial) and Vδ2+ (circulating), are defined based on the recombination of the TCR γ and δ chains. Vδ1+ T cells are predominantly located in peripheral tissues and possess robust anticancer capacities without the requirement for opsonizing antibodies, as Vδ2+ T cells do (*24, 25*). Adoptive transfer of polyclonal γδ T or Vδ1+ T cells leads to better survival of mice with ovarian cancer xenografts than adoptive transfer of Vδ2+ T cells (*26*). Nevertheless, the clinical use of Vδ1+ T cells has been fairly limited due to the scarcity of a reliable expansion protocol for selective and sustained proliferation of Vδ1+ T cells *ex vivo*. Instead, Vδ2+ T cells are used in most clinical trials because they can be easily expanded with amino bisphosphonates (e.g., zoledronic acid) or phosphoantigens. To conduct preclinical functional studies with Vδ1+ T cells for their therapeutic potential, our group utilizes a sequential cytokine stimulation protocol for *ex vivo* clinical-grade expansion of γδ T cells preferentially enriched for the Vδ1+ subset from the peripheral blood of healthy donors. Over the course of 21 days, the percentage of total γδ T cells in the CD3+ cell population increased from less than 10% to over 90%, while Vδ1+ T cells accounted for approximately 70% of all γδ T cells (**Fig. 2A and fig. S2**). To evaluate the immunophenotypes of expanded γδ T cells, we performed single-cell analysis on PBMCs at baseline and after *ex vivo* expansion using mass cytometry with an antibody panel designed to characterize T cell and MHC-unrestricted immunity-related receptors. We used t-distributed stochastic neighbor embedding (t-SNE) to investigate the differences and cellular heterogeneity within T cells before and after *ex vivo* expansion. First, preferential enrichment of the Vδ1+ subset from the peripheral blood after 14 days of expansion was noted (**Fig. 2B and fig. S3, A and B**). Within the expanded Vδ1+ subset, we observed differential expression of MHC-unrestricted immunity-related receptors, including NKG2D, CD226, CD244, NTB, CRACC, and NKp30, and the expression levels of these receptors were highly correlated (**Fig. 2C**). Most importantly, our *ex vivo* expanded γδ T cells express markers for cytolytic degranulation (e.g., CD107a) as well as secrete antitumor effector cytokines (i.e., TNF-α and IFN-γ) upon stimulation with pan-T cell activators — phorbol myristate acetate (PMA) and ionomycin. On the other hand, the expanded γδ T cells secrete almost no protumor or negative regulatory cytokines, such as IL-17A and IL-10 (**Fig. 2B and fig. S3B and S4**). The data indicate the antitumor phenotypes of these *ex vivo* expanded γδ T cells for potential therapeutic uses.

**Fig. 2.**
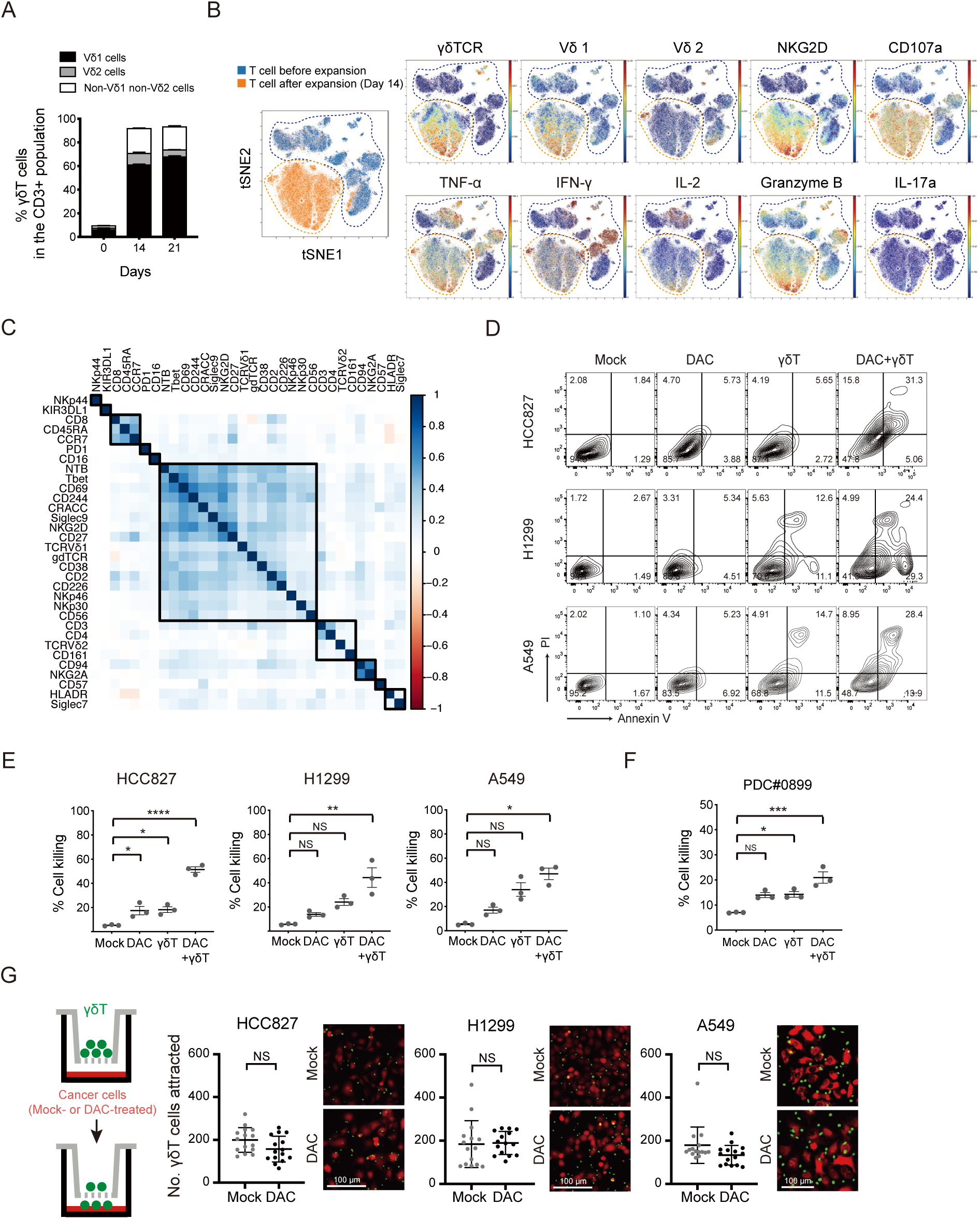
Decitabine (DAC) enhances γδ T cell-mediated cytolysis of lung cancer cells. (**A**) Bar graphs showing percentages of γδ T cell subsets (i.e., Vδ1, Vδ2, non-Vδ1 non-Vδ2 cells) in the CD3+ population from the peripheral blood of a healthy donor following *ex vivo* expansion at day 0, 14 and 21. Data are presented as mean ± standard error of the mean (SEM). (**B**) Overlay of tSNE maps of CD3+ T cells from PBMC at baseline and after *ex vivo* expansion analyzed by mass cytometry (CyTOF). Each dot represents a single cell. Cells enclosed in the blue or orange dashed lines are cells enriched at baseline or after expansion, respectively. For individual markers, the color represents the expression level of the indicated markers. Red is high, and blue is low. (**C**) Pearson correlation of markers expressed in *ex vivo* expanded γδ T cells analyzed by mass cytometry. The rectangles indicate groups with distinctive marker expression signatures according to the hierarchical clustering. (**D**) Representative flow cytometric analysis of Annexin V and propidium iodide (PI) apoptosis assays in human lung cancer cell lines — HCC827, H1299, A549 — upon treatments with 100 nM decitabine alone, γδ T cells alone or DAC/γδ T cell combination. The effector to target (E:T) ratio is 3:1. (**E**) Dot plots showing three biological replicates of Annexin V and propidium iodide (PI) apoptosis assays in human lung cancer cell lines described in (**D**). Data are presented as mean ± SEM. *p* value is calculated by one-way ANOVA with Tukey’s multiple comparison test (*, *p* < 0.05; **, *p* < 0.01; ***, *p* < 0.001; ****, *p* < 0.0001). (**F**) Annexin V and propidium iodide apoptosis assays of a patient-derived lung cancer cell line from malignant pleural effusion, PD#0899 (mean ± SEM, n = 3). (**G**) Transwell migration assays of γδ T cells (upper chamber) towards mock- or DAC-treated lung cancer cells (lower chamber). Numbers of γδ T cells in the lower chambers are counted per high power field and presented as mean ± standard errors in the dot plots. Representative images of γδ T cells (Hoechst 33342-labeled; green) and lung cancer cells (Calcein-retained; red) in the lower chambers are shown. Data are presented as mean ± standard deviation (SD). *p* value is calculated by the Mann-Whitney test. Scale bar: 100 μm.

### Decitabine enhances γδ T cell-mediated cytolysis of lung cancer cells

Subsequently, we investigated whether pretreatment with low-dose DAC may potentiate the susceptibility of lung cancer cells to attack by γδ T cells. First, we found that *in vitro* coculture of untreated lung cancer and γδ T cells at an effector to target (E:T) ratio of 3:1 caused 20-30% cytolysis of A549 human lung cancer cells (**fig. S5A**). As anticipated, 72-hour daily treatment with 100 nM DAC on human lung cancer cells (i.e., HCC827, H1299, and A549 cells) significantly potentiates the killing of lung cancer cells by γδ T cells at the same E:T ratio of 3:1. On the other hand, either treatment alone resulted in minimal or moderate cell death using Annexin V and propidium iodide apoptosis assays (**Fig. 2, D and E**). We observed similar potentiating effects in several other lung cancer cell lines, including H2981, PC9, PC9-IR (Iressa resistant), H157, CL1-0, and CL1-5, as well as patient-derived lung cancer cells from a malignant pleural effusion, PDC#0899 (**fig. S5B and Fig. 2F**). DAC’s potentiation effect on γδ T cell killing is also demonstrated using an electric cell-substrate impedance sensing (ECIS^TM^) system, a biophysical approach for real-time monitoring of the γδ T cell killing process. γδ T cell-mediated cytolysis usually takes place within 30 minutes to a few hours. The effect can be enhanced by DAC pretreatment at a dose that does not cause significant cytotoxicity (**fig. S5C**). Notably, this potentiating effect can also be observed in other tumor types, such as HCT116 colon cancer cells (**fig. S5D**).

Activated γδ T cells may trigger cancer death via the granule exocytosis-dependent cytotoxic pathway upon direct contact or through secretory TRAIL-mediated noncontact cytotoxicity that engages TRAIL receptors and the downstream apoptotic pathway (*27, 28*). To assess the significance of secretory TRAIL-mediated apoptosis in DAC-potentiated γδ T cell killing, we used a transwell system that allows for diffusion of γδ T cell-secreted TRAILs to reach cancer cells without direct cell contact. No significant cancer cell death was observed after 24 hours (**fig. S6**), suggesting that direct cell contact is required for DAC-potentiated γδ T cell killing. In addition, we evaluated whether DAC enhances γδ T cell chemotaxis using a transwell coculture system that allows γδ T cells to pass through the membrane, and there was no significant difference between the numbers of γδ T cells migrating toward mock-treated or DAC-treated lung cancer cells in the bottom wells. The data suggest that the potentiating effect of DAC is not dependent on increased chemoattraction of γδ T cells by DAC-primed cancer cells (**Fig. 2G**).

### Decitabine facilitates immune synapse formation between lung cancer and γδ T cells

Effective lysis of cancer cells by immune cells relies on functional immune synapses to facilitate directional and coordinated delivery of lytic granules (*29*). We investigated the efficiency of immune synapse formation between γδ T cells and DAC-pretreated cancer cells by immunofluorescence staining of phosphotyrosine (pTyr), a marker of immune synapses with active signaling (*30*). We observed a much higher number of immune synapses formed between γδ T and DAC-treated H1299 lung cancer cells than by mock-treated cells (**Fig. 3Α**). In search of the key molecules involved in the synaptic interaction, we compared the DAC-induced surface proteome of two human lung cancer cell lines — H1299 and A549, both of which showed enhanced γδ T cell killing following DAC treatment. Among the identified proteins, intercellular adhesion molecule 1 (ICAM-1) was highly upregulated in both cell lines (**Fig. 3B**). Western blotting confirmed that DAC can significantly upregulate ICAM-1 proteins in many lung cancer cell lines as well as a patient-derived lung cancer cell line from a malignant pleural effusion, PDC#062 (**Fig. 3C**). ICAM-1 is a surface glycoprotein and a member of the immunoglobulin superfamily. It is present in immune cells, endothelial cells, and epithelial cells as a general adhesion molecule. Adhesion of ICAM-1 on target cells to lymphocyte function-associated antigen-1 (LFA-1) on lymphocytes is known to initiate immune synapse formation, facilitate αβ T cell activation and mediate the delivery of cytotoxic granules from conventional αβ T cells toward target cells (*31*). In the case of γδ Τ cell immune synapses, we demonstrated that ICAM-1 is localized within the immune synapse along with other classical immune synapse proteins, including LFA-1 and linker for activation of T cells (LAT) (**Fig. 3D**).

**Fig. 3.**
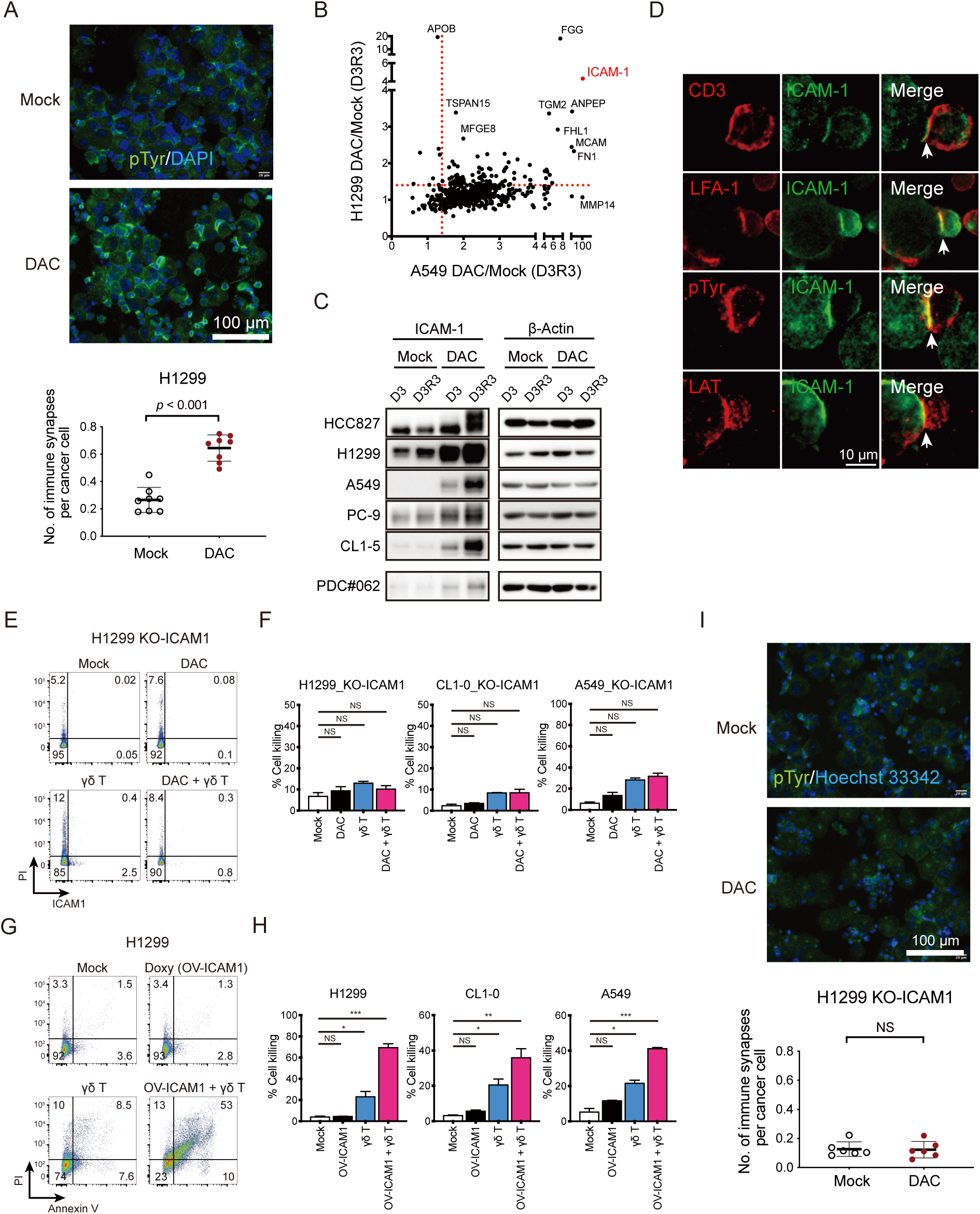
Decitabine (DAC) facilitates immune synapse formation between lung cancer and γδ T cells. (**A**) Immunofluorescence imaging of immune synapses between H1299 lung cancer cells and γδ T cells by phosphotyrosine (pTyr) staining. H1299 lung cancer cells are pretreated with phosphate-*buffered* saline (PBS) or DAC prior to coculture with γδ T cells. Quantifications of immune synapses per cancer cell on eight randomly taken high power fields for each treatment are shown in the dot plots (mean ± SD). Scale bar: 100 μm. *p* value is calculated by the Mann-Whitney test. (**B**) A scatter plot of DAC-induced surface proteomes in H1299 (y-axis) and A549 (x-axis) human lung cancer cells following daily treatment of 100 nM DAC for 72 hours and culture in drug-free medium for 3 days (D3R3). ICAM-1 is among the top upregulated surface proteins by DAC in both cells. (**C**) Western blot analyses of ICAM-1 protein expression in mock-treated vs. DAC-treated human lung cancer cells. D3: daily treatment of 100 nM decitabine for 72 hours. D3R3: daily treatment for 72 hours, followed by a 3-day rest period in drug-free medium. β-actin: loading control. (**D**) Immunofluorescence staining of ICAM-1 and immune synapse molecules (e.g., LFA-1, LAT) at immune synapses formed between γδ T cells and DAC-treated H1299 lung cancer cells. Scale bar: 10 μm. (**E**) Representative flow cytometric dot plot showing H1299 lung cancer cells with CRISPR-knockout of ICAM1 (KO-ICAM1) subject to γδ T cell killing for 2 hours. The effector to target (E: T) ratio is 3:1. Lung cancer cells are pre-treated with mock, DAC alone, γδ T cells alone or a combination of DAC and γδ T cells. The X-axis denotes surface ICAM1 levels. Y-axis represents signal intensities of propidium iodide. (**F**) Bar graphs showing percent cell death of human lung cancer cell lines (i.e., H1299, CL1-0, and A549) with CRISPR-knockout of ICAM-1 subject to γδ T cell killing for 2 hours. Cell death is measured by Annexin V and propidium iodide apoptosis assays (mean ± SEM, n = 3). Statistical significance is determined by one-way ANOVA test. (**G**) Representative flow cytometric dot plot showing H1299 lung cancer cells with a Tet-on expression system of ICAM1 (OV-ICAM1) subject to γδ T cell killing for 2 hours. Doxycycline (1 μg/mL) is added 24 hours prior to coculture to induce ICAM-1 protein expression. Cell death is measured by Annexin V (x-axis) and propidium iodide (y-axis) apoptosis assays. (**H**) Bar graphs showing cell death of human lung cancer cell lines (i.e., H1299, CL1-0, and A549) with ICAM-1 over-expression subject to γδ T cell killing for 2 hours. E:T ratio is 3:1. Cell death is measured by Annexin V and propidium iodide apoptosis assays. Statistical significance is determined by one-way ANOVA test (\**p* < 0.05, \*\**p* < 0.01, ***, *p* < 0.001). (**I**) Immunofluorescence imaging of immune synapses between H1299 KO-ICAM1 lung cancer cells and γδ T cells by phosphotyrosine (pTyr) staining. Scale bar: 100 μm. Quantifications of immune synapses per cancer cell on six randomly taken high power fields for each treatment are shown in the dot plots (mean ± SD). *p* value is calculated by the Mann-Whitney test.

Notably, knockout of ICAM-1 (KO-ICAM1) with CRISPR technology (**figs. S7 and S8**) in lung cancer cells (i.e., A549, H1299, and CL1-0) ultimately diminished the potentiation effects of DAC for γδ T cell killing in all three cell lines tested (**Fig. 3, E and F**). On the other hand, overexpression of ICAM-1 (OV-ICAM1) in lung cancer cell lines with a Tet-On system (**fig. S8**) markedly enhanced γδ T cell-mediated cytotoxicity on human lung cancer cells, mimicking the effect of DAC pretreatment (**Fig. 3, G and H**). As anticipated, DAC treatment of ICAM-1-knockout cells failed to enhance immune synapse formation between lung cancer and γδ T cells (**Fig. 3I**). In contrast, other immune synaptic proteins, such as putative ligands for γδ TCR, including CD1d complexes (*32*), BTN3A (*33*), BTNL3 (*34*) and others (*35*), are not consistently upregulated by DAC in all the cancer cell lines from our surface proteomics study (**fig. S9**) and may not be essential for DAC’s potentiating effect on γδ T cell cytotoxicity. Collectively, the data suggest that ICAM-1-mediated adhesion and stabilization of the synaptic structure is critical for DAC’s potentiating effects on γδ T cell antitumor immunity.

### Decitabine stabilizes the immune synaptic cleft to facilitate tumor lysis via strengthening the actin cytoskeleton

Classical immune synapses centered around the TCR/MHC complex are specialized structures formed between T cells and antigen-presenting cells (APCs) or T cells and target cells to facilitate vesicular traffic-mediated intercellular communication or tumor lysis (*31, 36*). As we further deciphered how DAC affects the synaptic structure to exert MHC-unrestricted cytotoxicity, we observed a marked accumulation of filamentous actin (F-actin) at the cancer cell membrane near the region of immune synapses (**Fig. 4, A and B**). When we knocked out ICAM-1 in H1299 lung cancer cells, there was a substantial decrease in DAC-induced F-actin accumulation at immune synapses (**Fig. 4C and fig. S10**), which was associated with a significant reduction in synaptic cleft width (**Fig. 4D**). In contrast, DAC-treated lung cancer cells with normal ICAM-1 expression established immune synapses with a prominent synaptic cleft between cancer and γδ T cells (**Fig. 4D**). Notably, pharmacological disruption of actin polymerization with cytochalasin B (Cyto B) abolished DAC-induced F-actin clustering and pTyr signaling at immune synapses but appeared to have minimal effects on ICAM-1 expression (**Fig. 4E and fig. S11A**). Additionally, cytochalasin B also diminished the DAC effects on the width of the synaptic cleft (**fig. S11, B and C**). Furthermore, cytochalasin B significantly counteracted DAC’s enhancing effects on immune synapse formation (**Fig. 4, F and G**). F-actin accumulation and larger sizes of the synaptic clefts are characteristics of activating instead of inhibitory immune synapses (*37*). Our data suggest that DAC remodels the actin cytoskeleton to facilitate the formation of activating immune synapses between cancer and γδ T cells.

**Fig. 4.**
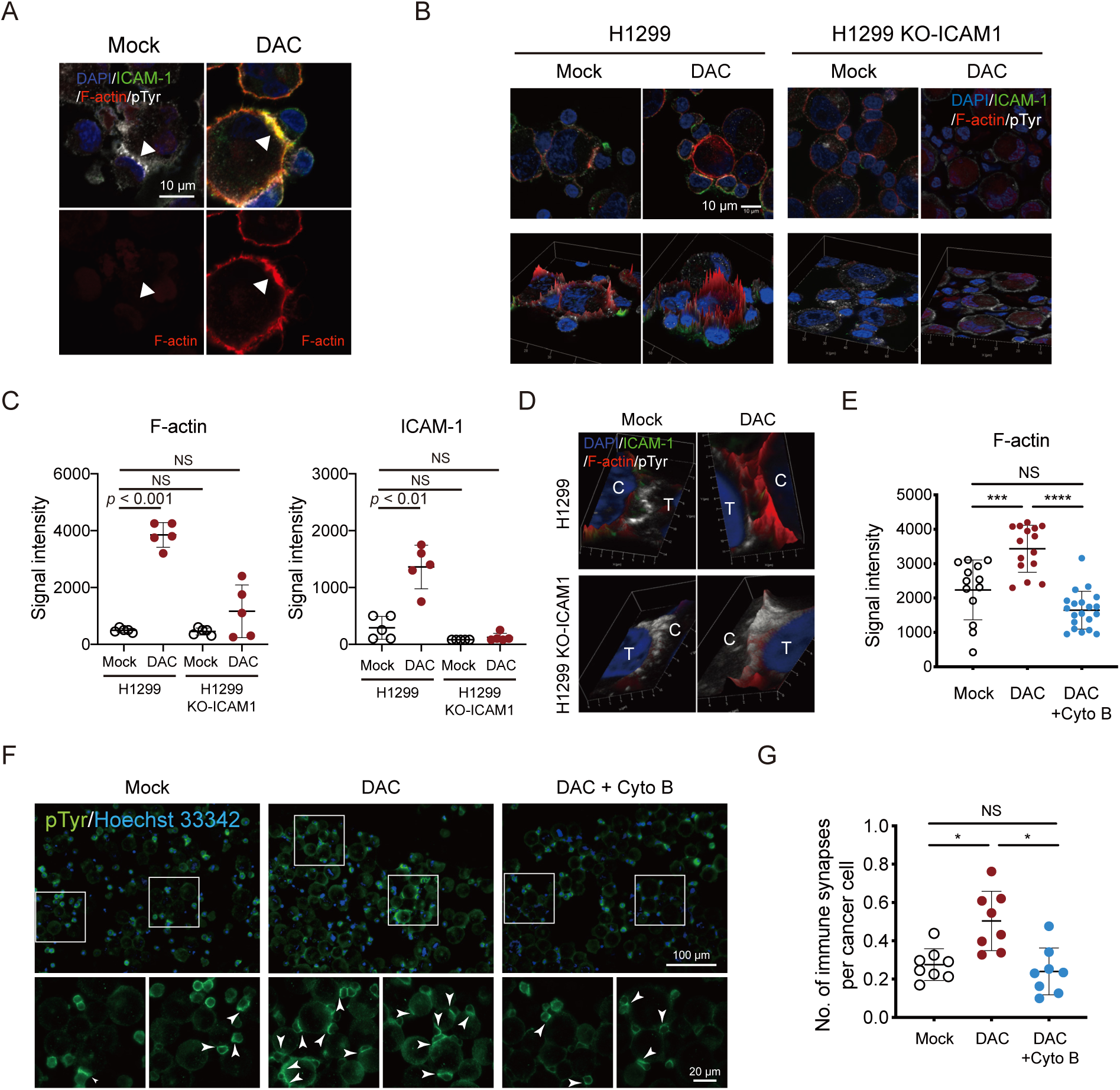
Decitabine stabilizes the immune synaptic cleft to facilitate tumor lysis via strengthening the actin cytoskeleton. (**A**) Immunofluorescence staining of F-actin (red), ICAM-1 (green), pTyr (phosphotyrosine, white) at immune synapses between γδ T cells and DAC-pretreated H1299 lung cancer cells at D3R3. Accumulation of F-actin beneath the cell membrane is noted in DAC-pretreated lung cancer cells. DAPI: 4′,6-diamidino-2-phenylindole, as a nuclear counterstain. Scale bar: 10 μm. (**B**) Representative immunofluorescence images of the interfaces between γδ T cells and H1299 lung cancer cells (parental vs. ICAM-1 knockout (KO-ICAM1)). Signals of F-actin (red) in the periphery of H1299 cancer cells are shown in two- and-a-half-dimensional (2.5D) images in the lower panels. Scale bar: 10 μm. (**C**) Dot plots of signal intensities of F-actin (left panel) and ICAM-1 (right panel) from five pTry-positive immune synapses between γδ T cells and H1299 lung cancer cells (parental or KO-ICAM1). *p* value is calculated by two-way ANOVA test. (**D**) Immunofluorescence images of immune synapses between γδ T cells (marked with T) and H1299 lung cancer cells (marked with C) stained for ICAM-1 (green), F-actin (Red) and phosphotyrosine (pTyr, white). Lung cancer cells (parental or KO-ICAM1) are pretreated with PBS (Mock) or 100 nM DAC and cocultured with γδ T cells at D3R3. (**E**) Dot plots of F-actin signal intensities at immune synapses between γδ T cells and H1299 cells. H1299 cells are pretreated with PBS (Mock), DAC alone or combination of DAC pretreatment (D3R3) and 1 μg/mL Cyto B (cytochalasin B, an inhibitor of actin filament polymerization) for 1.5 hours prior to coculture with γδ T cells (mean ± SD). *p* value is calculated by one-way ANOVA with Tukey’s multiple comparisons test (***, *p* < 0.001; ****, *p* < 0.0001). (**F**) Representative immunofluorescence images of immune synapses (pTyr staining) between γδ T and H1299 cells pretreated with PBS (Mock), DAC alone, and combination of DAC and Cyto B. Blow-up images of the square areas for each treatment are shown in the lower panels. Arrows denote immune synapses between γδ T and H1299 cells. Scale bar: 100 μm (upper) and 20 μm (lower panels). (**G**) Dot plots showing numbers of immune synapses per cancer cell on eight randomly taken high power fields for H1299 cells pretreated with PBS (Mock), DAC, and combination of DAC and Cyto B (mean ± SD). *p* value is calculated by one-way ANOVA with Tukey’s multiple comparisons test (*, statistical significance).

### Depletion of DNMTs induces γδ T-sensitive cytoskeletal gene patterns at the cancer side of the immune synapse

The cytoskeleton plays an essential role in enabling cell motility, maintaining cell shape, and directing axon guidance. It also serves as an integral part of neurologic or immune synapses (*38, 39*). While it has been known that proper cytoskeleton dynamics and arrangements in immune cells are critical for satisfactory immune responses, the expression patterns of the cytoskeleton at the cancer cell side of the immune synapse, their regulation, and the associated functional consequences remain incompletely understood. In light of our observation of DAC-induced actin cytoskeleton reorganization (**Fig. 4**), we speculate that there may be coordinated regulation of immune-related cytoskeletal gene networks by disruption of DNMTs. Indeed, genome-wide mRNA-seq data of five lung cancer cell lines (i.e., A549, CL1-0, CL1-5, PC9, and H1299) after DAC treatment showed a significant induction of genes/enzymes involved in cytoskeletal dynamics and reorganization, including CORO1A (*40*), HCLS1 (*41*), FES (*42*), among others (**Fig. 5A**). Gene set enrichment analysis (GSEA) also revealed a striking enrichment for gene sets related to actin cytoskeleton reorganization as well as intermediate filament-based processes. The enrichment was highly associated with the upregulation of ICAM-1. In contrast, the gene sets related to microtubules appeared to be downregulated (**Fig. 5, B and C**). Likewise, genetic depletion of DNMTs recapitulates similar cytoskeletal gene expression profiles, as we analyzed the transcriptomic data of colon cancer cell lines (e.g., HCT116 and DLD1) subject to shRNA targeting of DNMTs (*11*) (**Fig. 5D**). In addition, the regulation of these cytoskeletal genes after DAC treatment appears to be a highly coordinated epigenetic process. When we examined promoter DNA methylation status of the genes in the three cytoskeletal gene modules — actin cytoskeleton, intermediate filaments and microtubules — in human lung cancer cells with Infinium MethylationEPIC BeadChips, we found that genes in the actin gene module had a higher promoter DNA methylation at baseline and became demethylated with DAC treatment. In contrast, genes in the microtubule module tended to have low baseline methylation levels, which were minimally altered by DAC (**Fig. 5E**).

**Fig. 5.**
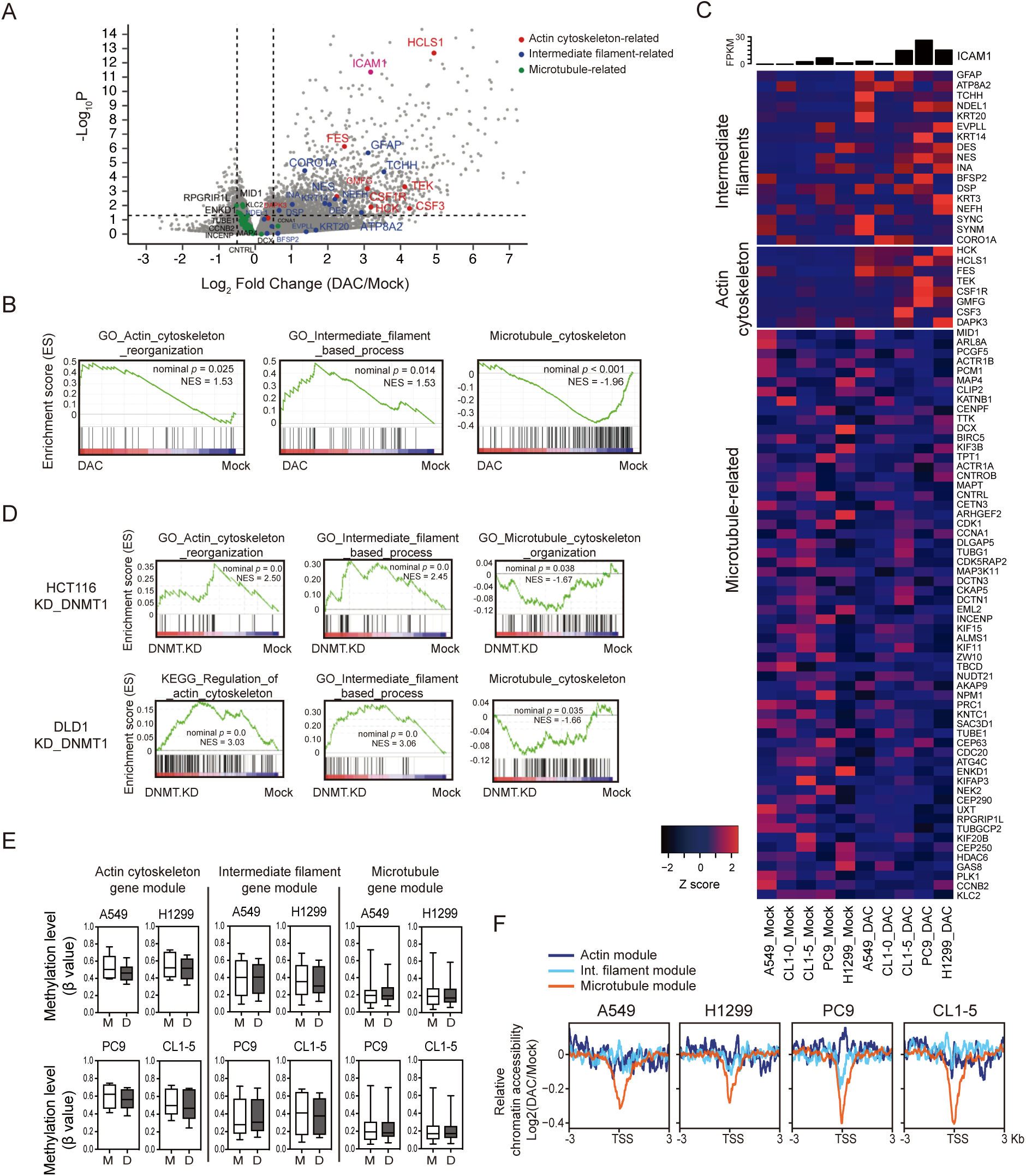
Depletion of DNMTs induces γδ T-sensitive cytoskeletal gene patterns at the cancer side of the immune synapse. (**A**) A volcano plot showing differentially expressed genes in DAC-treated vs. Mock-treated human lung cancer cells (i.e., A549, CL1-0, CL1-5, PC9, and H1299). The y-axis denotes statistical significance (-log10 of *p-value*), and the x-axis displays the log2 fold change values between the DAC-treated and the Mock-treated groups. Genes related to the actin cytoskeleton, intermediate filaments, microtubule are marked in red, blue and green, respectively. (**B**) Gene Set Enrichment Analysis (GSEA) of mRNA-seq data in five DAC-treated lung cancer cell lines — A549, CL1-0, CL1-5, PC9, and H1299. Gene sets related to actin-cytoskeleton reorganization, intermediate filament-based process, and microtubule-related gene modules are shown. NES: Normalized Enrichment Score. (**C**) Heatmap showing mRNA expression of core enrichment genes for actin-cytoskeleton, intermediate filament, and microtubule-related processes in DAC-treated and mock-treated human lung cancer cells measured by mRNA-seq. (**D**) Gene Set Enrichment Analysis (GSEA) of mRNA-seq data in HCT116 and DLD1 human colorectal cancer cell lines subject to shRNA knockdown of DNA methyltransferase 1 (DNMT1). Gene sets related to actin-cytoskeleton reorganization, intermediate filament-based process, and microtubule-related gene modules are shown. NES: Normalized Enrichment Score. (**E**) Box plots with median and 95% confidence interval showing promoter methylation status of genes in the actin cytoskeleton-, intermediate filament-, and microtubule-related gene modules in human lung cancer cells treated with 100 nM DAC for three days followed by a 3-day drug-free culture (D3R3). Methylation data are analyzed by Infinium MethylationEPIC arrays. (**F**) Relative chromatin accessibility around TSS of genes (−3 to +3 kb) in the actin-cytoskeleton reorganization, intermediate filament-based process, and microtubule-related modules in DAC-treated vs. mock-treated lung cancer cell lines.

Moreover, emerging evidence has suggested that DAC may alter epigenetic regulatory circuits, in addition to its well-known DNA-demethylating properties, to exert multifaceted transcriptional control of a gene (*43*). To investigate whether and how chromatin accessibility may be involved in DAC transcriptional regulation, we performed Omni-ATAC-seq (*44*) in these human lung cancer cells after DAC treatment (**fig. S12A**). For overall pattern exploration, we stratified the DAC-upregulated genes into three groups based on their basal promoter methylation status (β > 0.6, 0.3 < β < 0.6, and β < 0.3) (**Fig. 12B**) and analyzed the changes in chromatin accessibility for each gene group before and after DAC treatment. Consistent with common speculation, chromatin accessibility at the transcription start site (TSS) inversely correlated with promoter DNA methylation levels. In particular, many upregulated genes with low basal promoter DNA methylation have preexisting accessible chromatin at the TSS, which remains accessible following DAC treatment (**fig. S12, B and C**). On the other hand, genes with high promoter DNA methylation level at baseline tended to have inaccessible chromatin, which gained modest but critical accessibility after DAC treatment (**fig. S12, B and C**). Moreover, peak calling by MACS2 (*45*) revealed more peaks of accessible chromatin at TSSs in DAC-treated lung cancer cells, consistent with extensive gene upregulation after DAC treatment (**fig. S12D**). Strikingly, a detailed examination of cytoskeletal gene modules shows a marked decrease in open chromatin regions at gene promoters of the microtubule-related genes downregulated by DAC (**Fig. 5F**). In contrast, moderate chromatin changes were observed at promoters of genes in actin- or intermediate filament-related processes (**Fig. 5F**). These data suggest that there are distinct patterns of DAC-mediated regulatory mechanisms for genes with different promoter DNA methylation levels. And preexisting chromatin accessibility at unmethylated promoters may predispose to DAC-induced gene upregulation.

### Network analysis reveals TP53 as a hub for interactions of immune synaptic molecules, the cytoskeleton and epigenetic proteins

Subsequently, we examined the epigenetic regulatory pattern of individual genes in each cytoskeletal module, such as DAPK3 (a serine/threonine kinase involved in actin cytoskeleton reorganization), EVPLL (a crosslinker of intermediate filament) and TUBE1 (tubulin epsilon chain). Consistent with our genome-wide analysis, DAC-induced upregulation of EVPLL was associated with DNA demethylation; on the other hand, DAC-induced downregulation of TUBE1 mainly correlated with decreased chromatin accessibility (**Fig. 6A**). Interestingly, we observed a mixed pattern of *ICAM1* promoter DNA methylation at baseline in multiple lung cancer cell lines despite the prevailing trend of increased mRNA expression after DAC treatment (**Fig. 6B**). In A549 and H1299 cells, the promoter region of *ICAM1* (approximately −500 to 0 bp) is DNA hypermethylated and became demethylated upon DAC treatment, which was associated with an increase in chromatin accessibility around the promoter CpG island (**Fig. 6C**). For another two cell lines, PC9 and CL1-5, the *ICAM1* promoter appears to be unmethylated and was unaltered by DAC treatment despite the increases in mRNA expression. Interestingly, DAC-induced chromatin accessibility alterations were modest, especially in PC9, where the chromatin at the *ICAM1* promoter was already highly accessible before drug treatment (**Fig. 6C**). These changes were also experimentally validated by locus-specific quantitative ATAC-PCR at the *ICAM1* promoter (**Fig. 6D**). We reason that binding of transcription factors to the open chromatin region may be required to trigger profound transcriptional activation of *ICAM1*. Using ChIP-seq data via the ENCODE portal (*46*), we identified 230 proteins that may bind to the *ICAM1* promoter in all available cell types. Among 230 proteins, RELB, NFKB2, STATS, and RUNX are located in the experimentally determined open chromatin regions at the *ICAM1* promoter in DAC-treated PC9 and CL1-5 cells and other lung cancer cell lines (**fig. S13A**). These transcription factors appear to respond to DNA demethylation and be upregulated by DAC (**fig. S13B**), which indicates a transactivating effect on the accessible *ICAM1* promoter. The data support a multilayered modulation of DAC on gene transcription in a context-dependent manner.

**Fig. 6.**
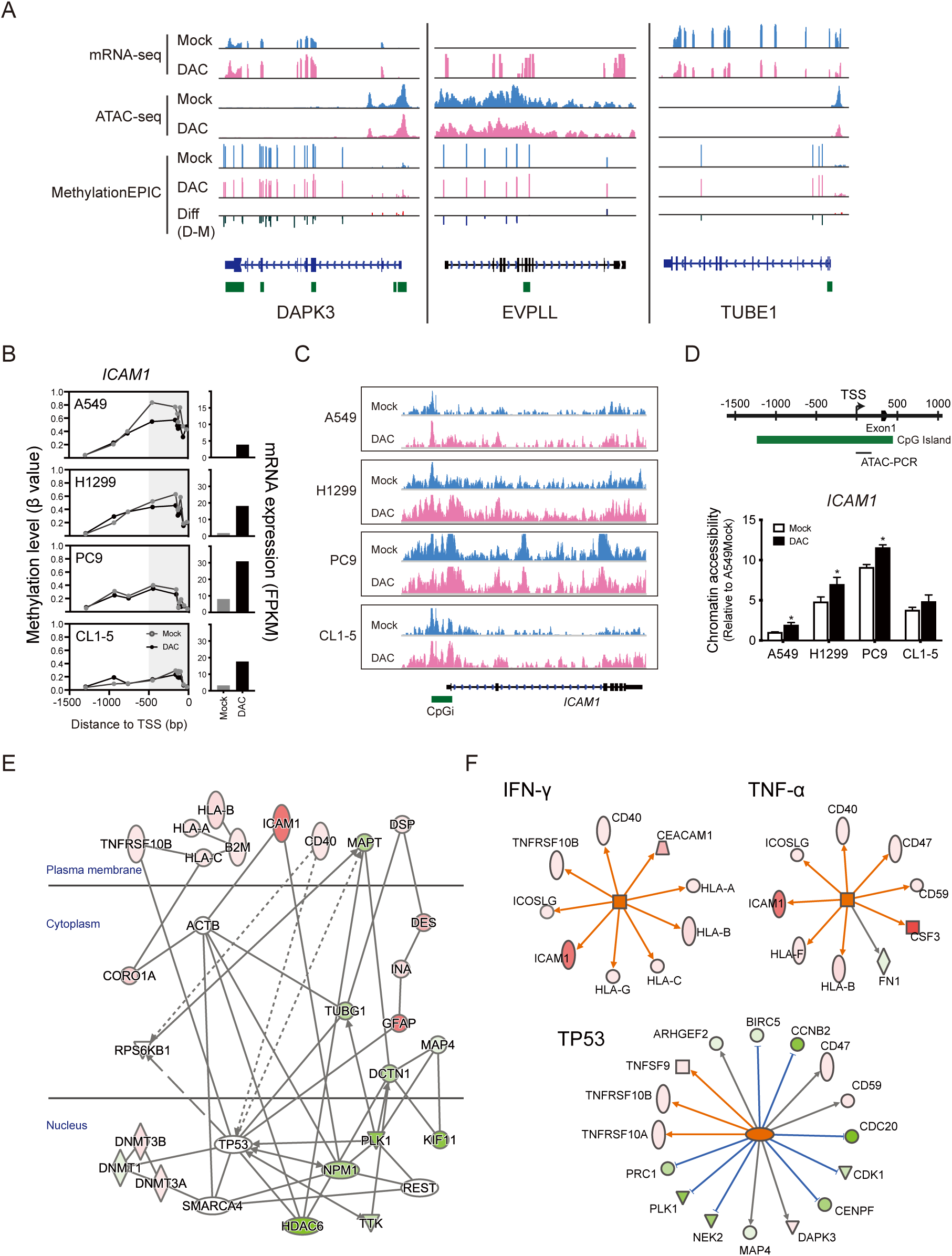
Network analysis reveals TP53 as a hub for interactions of immune synaptic molecules, the cytoskeleton and epigenetic proteins. (**A**) Visualization of multi-omics data (i.e., mRNA-seq, Omni-ATAC-seq, and MethylationEPIC arrays) for *DAPK3*, *EVPLL,* and *TUBE1* in H1299 lung cancer cells. (**B**) Promoter methylation status and mRNA expression levels of the *ICAM1* gene measured by Infinium MethylationEPIC arrays (left panels) and mRNA-seq (right panels) in human lung cancer cells treated without and with DAC 100 nM DAC for 3 days followed by a 3-day drug-free culture. (**C**) Open chromatin regions in the promoter areas of the *ICAM1* gene in human lung cancer cells upon 100 nM DAC treatment analyzed by Omni-ATAC-seq. The green bar represents a CpG island. (**D**) Validation of Omni-ATAC-seq by quantitative real-time PCR on transposase-accessible chromatin at the *ICAM1* promoter of human lung cancer cells subject to daily treatment of 100 nM DAC treatment for 3 days, followed by a 3-day drug-free culture. Experiments are performed in triplicates, and data are presented as mean ± SD. *p* value was calculated by unpaired t test (*, p < 0.05). (**E**) IPA Network analysis of mRNA expression changes in human lung cancer cells treated by DAC reveals coordinated changes of the immune-related surface molecules and the cytoskeleton-associated genes. (**F**) IPA upstream regulator analysis of mRNA expression changes in human lung cancer cells treated by DAC. T cell effector cytokines such as TNF-α and IFN-γ may enhance DAC-induced expression changes of immune-related molecules and ICAM-1 in lung cancer cells. TP53 is a potential master regulator for cancer cytoskeleton reorganization essential for DAC-potentiated γδ T cell killing.

Notably, gene network analysis of DAC transcriptomes in lung cancer cell lines by Ingenuity Pathway Analysis (IPA®) revealed an intimate interconnectedness between immune synaptic molecules, the cytoskeleton, and epigenetic proteins (**Fig. 6E**). As shown in the network, we found a general downregulation of genes involved in microtubule organization (i.e., TUBG1, DCTN1, and PLK1). In contrast, genes participating in actin and intermediate filament dynamics (i.e., CORO1A, GFAP, and DES) were upregulated. *ICAM1* bridges surface immune receptors/HLA molecules to the cytoskeleton in the cytoplasm, which links to TP53 and other epigenetic modifiers (i.e., DNMTs, HDACs, and SMARCA4) in the nucleus (**Fig. 6E**). Further upstream regulator analysis showed that T cell effector cytokines, such as IFN-γ and TNF-α, may enhance the DAC-induced expression pattern of immune surface molecules, including ICAM-1, ICOSLG, and HLAs (**Fig. 6F**). In addition, TP53 appears to be a hub gene mediating the coordinated changes of immune surface molecules the cytoskeleton in response to epigenetic modifications, implying that a functional TP53 network is necessary for effective immune potentiating effects by DAC (**Fig. 6F**).

### Single-cell mass cytometry characterizes functional γδ T cell subsets modulated by decitabine

To translate the *in vitro* finding of DAC’s potentiating effects on γδ T cell killing into potential therapeutic efficacy, we first evaluated how DAC affects γδ T cells since the drug’s effects on γδ T cells are incompletely understood. Again, we used mass cytometry to profile both phenotypic and functional immune parameters of expanded γδ T cells with or without 10 nM DAC treatment. Data from both groups of cells were clustered together by an x-shift algorithm, and the frequency of each subpopulation in untreated and DAC-treated expanded γδ T cells was calculated. As shown in **Fig. 7A**, 14 clusters within CD3+ T cells were revealed (termed T1 through T14, ranked by the frequency differences between the untreated and DAC-treated groups), each with distinct phenotypic and functional effector signatures. Importantly, the top two cell clusters induced by DAC treatment correspond to Vδ1 and express higher levels of CD226, CD244, CD2, and CRACC together with stronger functional effectors, including CD107A, TNF-α, granzyme B, IL-2 and IFN-γ, than those expressed by the clusters decreased in cell frequency after DAC treatment (**Fig. 7B**). The result was corroborated by Vδ1 T cells expanded from another five healthy individuals treated with 10 nM DAC. We observed a trend of increased production of antitumor effector cytokines, such as IFN-γ and TNF-α (**Fig. 7C**). Moreover, following DAC treatment, there was a marked increase in the percentages of polyfunctional Vδ1+ cells that coexpressed two or more effector cytokines in three of the five individuals (donors #1, #3 and #5) (**Fig. 7D**). As polyfunctional T cells are often considered the hallmark of protective immunity (*47–49*), the data further strengthen the rationale of combination therapy using DAC and adoptive transfer of γδ T cells.

**Fig. 7.**
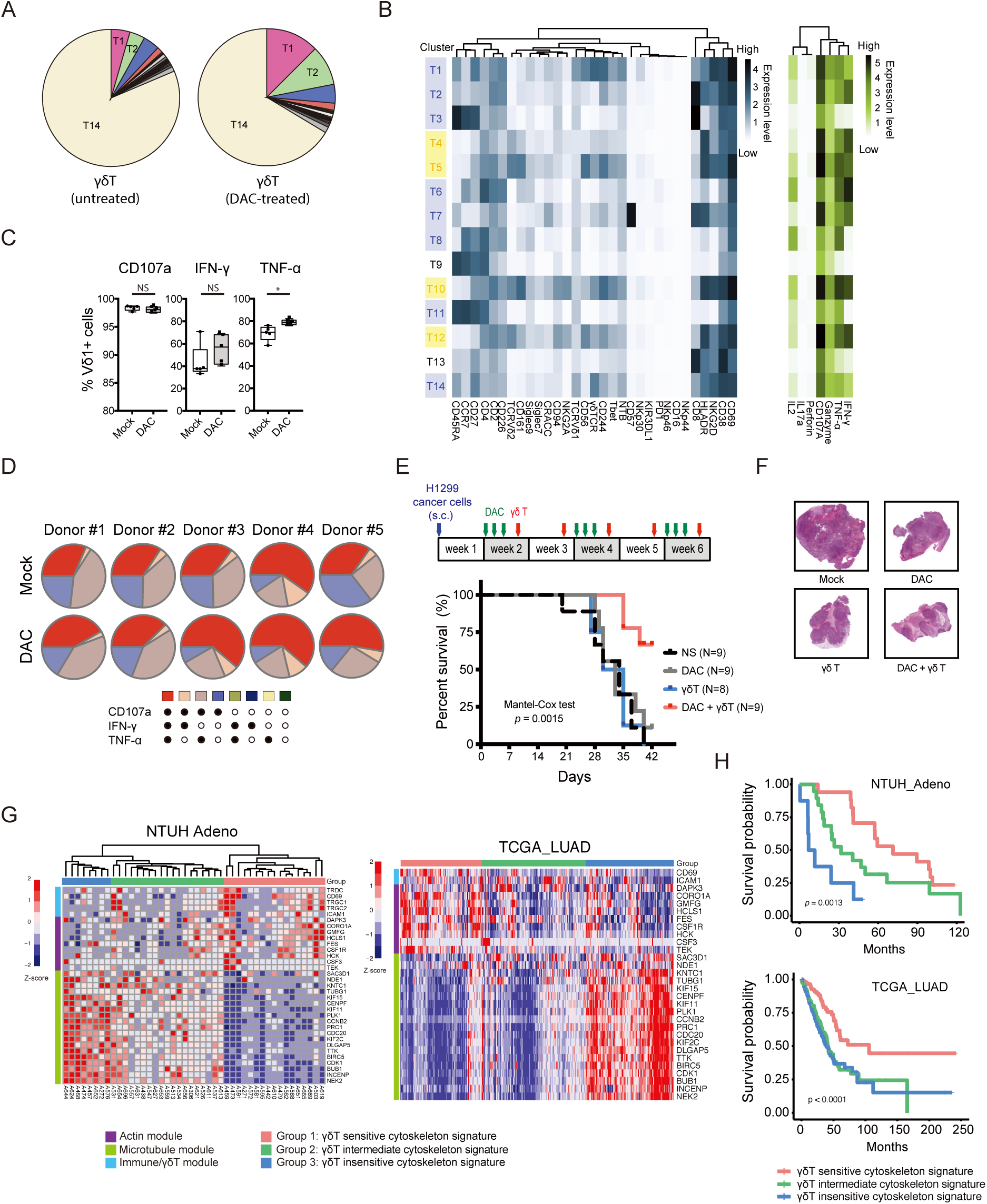
Combination therapy of decitabine (DAC) and γδ T cells prolongs survival in mice bearing lung cancer xenografts. (**A**) Pie charts showing the cell frequencies of 14 cell clusters (T1 to T14) revealed by clustering analysis of *ex vivo* expanded γδ T cells with or without DAC treatment based on 38 mass cytometric markers. (**B**) Heatmaps showing the mean intensity of each marker in the 14 cell clusters (T1 to T14). Dark blue and dark green are high, and white is low. Clusters belong to Vδ1 or Vδ2 are labeled in blue and yellow, respectively. (**C**) Flow cytometric analysis of effector cytokine production by *ex vivo* expanded γδ T cells from 5 healthy donors at D3R3 with 10 nM DAC or mock treatment. Data is presented in box and whisker plot showing the range minimum to a maximum of the percentages of Vδ1+ γδ T cells expressing individual cytokines. *p* value is calculated by the Mann-Whitney test (*, *p* < 0.05). (**D**) Pie charts showing percentages of monofunctional and polyfunctional *ex vivo* expanded Vδ1+ γδ T cells from 5 healthy donors at D3R3 with 10 nM DAC or mock treatment. Percentages of Vδ1+ γδ T with a concurrent expression of three effector cytokines are marked in red. (**E**) The *in vivo* experiment of NOD-scid IL2rg^null^ (NSG) mice bearing H1299 human lung cancer xenografts treated by DAC, adoptive human γδ T transfer, or both. For every cycle of drug treatment, DAC is administered intraperitoneally for three consecutive days, followed by intravenous injection of *ex vivo* expanded γδ T cells at day 5 and day 12. Kaplan-Meier survival curve of NSG mice in each treatment group is shown. *p* value is calculated by the Mantel-Cox test. (**F**) Ηematoxylin and eosin (H&E) staining of representative mouse tumors in each treatment group. Images of the whole tumor are generated by a digital slide scanner. (**G**) A heatmap of immune cytoskeleton gene signature derived from mRNA-seq data of primary lung adenocarcinoma tumor tissues in patients at National Taiwan University Hospital (NTUH, *left*) and from The Cancer Genome Atlas (TCGA, *right*). (**H**) Overall survival analysis of the NTUH (*upper*) and TCGA (*lower*) lung adenocarcinoma patient cohorts stratified by immune cytoskeleton gene signatures associated with different γδ T susceptibilities. *p* value is calculated by the Mantel-Cox test.

### Combination therapy of DAC and adoptive transfer of γδ T cells prolongs survival in mice bearing lung cancer xenografts

Subsequently, we investigated the effects of combination therapy with DAC (intraperitoneal injection) followed by the adoptive transfer of *ex vivo* expanded human γδ T cells (intravenous injection) in immunocompromised NSG^TM^ (NOD.Cg-Prkdc^scid^ Il2rg^tm1Wjl^/SzJ) mice bearing H1299 human lung cancer xenografts (**Fig. 7E**). The mice in the combination therapy group had significantly better overall survival than the mice receiving normal saline or subject to either treatment alone **(Fig. 7E**). Pathologically, the tumor tissues in the combination group were not only smaller but also displayed loose architecture and marked fibrosis between small nodules. In contrast, tumor tissues in the control group appeared to be hypercellular, with regions of hemorrhage and necrosis **(Fig. 7F**). The data demonstrate the promise of combining DAC with the adoptive transfer of γδ T cells in treating lung cancer.

### Cytoskeletal gene signatures correlate with overall survival in lung adenocarcinoma patients

Finally, we examined the RNA-seq data from the primary lung cancer tissues at the National Taiwan University Hospital and the TCGA portal to determine the clinical significance of DAC-induced cytoskeleton signatures. Unsupervised clustering of primary lung adenocarcinoma tissues based on the core enrichment genes from the actin-, intermediate filament-, and microtubule-related gene modules (**Fig. 5C**) revealed three groups: γδ T cell-sensitive, γδ T cell-intermediate, and γδ T cell-insensitive (**Fig. 7G**). Remarkably, we observed the best overall survival in the patients with the γδ T cell-sensitive signature (the actin and intermediate filament modules upregulated and microtubule module downregulated) (**Fig. 7H**). In contrast, the γδ T cell-insensitive signature was associated with the worst prognosis in both cohorts (**Fig. 7H**). The data highlight the importance of epigenetic reorganization of the cytoskeleton as a clinical indicator for patient outcome and may be used for patient selection to maximize the benefits of γδ T cell-based therapy in lung cancer patients.

## DISCUSSION

DNA methyltransferase inhibitors are versatile immunomodulators (*3, 12*), aside from their antitumor effects (*50*). These drugs not only may enhance the immunogenicity of cancer cells (*4, 7, 8, 51*) but also have been shown to modulate T cell fate decisions, differentiation, and exhaustion (*9, 10*). Using a quantitative surface proteomics approach combined with epigenomic/transcriptomic analysis, we uncovered an immunomodulatory effect of DNMTis to facilitate tumor cytolysis by MHC-unrestricted γδ T cell-based therapy via upregulation of adhesion molecules and reorganization of the immunosynaptic cytoskeleton networks to strengthen the encounter between cancer and γδ T cells.

γδ T cells are a distinct subset of T cells that combine adaptive characteristics and rapid, innate-like responses. These unconventional T cells sense molecular stress signals from pathogen-infected or stressed cells and display broad tumor-targeting capabilities in an MHC-independent manner (*52*). Earlier clinical trials of adoptive autologous or allogeneic γδ T cell transfer demonstrated encouraging therapeutic responses in various cancer types; nevertheless, there is still room for improvement in clinical outcomes (*23, 53*). Our team developed a sequential cytokine stimulation protocol for selective expansion of Vδ1+ cells, the dominant γδ T cell subtype residing in the tissues, and a highly potent antitumor effector without the need for opsonizing antibodies in the tumor lytic process (*24*). γδ T cells are composed of heterogeneous subsets with pleiotropic immune effector functions (*22*). Our expansion protocol enriches γδ T cells with antitumor immunity instead of the protumor IL-17-producing γδ T subsets that have been shown to promote tumor progression (*54, 55*). This clinical-grade expansion protocol for Vδ1+ cells may facilitate the refinement of current γδ T cell therapy, as well as the development of dual-functional chimeric antigen receptor-engineered human γδ T cells (CAR-γδ T cells) that respond to both stress signals and specific cancer antigens (*56, 57*).

Moreover, our data demonstrate DAC’s potentiation effects on lung cancer cells for γδ T cell killing. This phenomenon is also observed in HCT116 colon cancer cells, which suggests that the combined use of epigenetic and adoptive γδ T cell therapy may be applicable in more than one cancer type. Interestingly, these immune-potentiating effects can be linked to epigenetic reprogramming of the immune synaptic-cytoskeletal networks. In fact, optimal orchestration of cytoskeletal networks in T cells is critical for proper organization of immune synaptic components, directional delivery of lytic granules, and transduction of TCR-activating signals (*58, 59*). On the other hand, the cytoskeletal organization on the target cell side of the immune synapse has been less explored and may be equally important for cytolytic activity upon immune attack (*38, 60*). Here, we show that the immune-related cytoskeleton pattern can be modulated by epigenetic therapy via coordinated gene network regulation as the result of DNA demethylation, as well as the ripple effects of DAC-induced DNMT degradation on chromatin modifiers or nucleosome remodelers. We also show that inhibiting actin polymerization significantly diminishes DAC’s potentiating effects on γδ T cell killing. The possibility of epigenetically converting an immune-resistant cytoskeleton pattern into an immune-sensitive pattern may provide a potential therapeutic strategy to maximize the clinical benefits of cell-based immunotherapy.

In addition, our holistic proteomics approach of characterizing surface immunome of lung cancer cells reveals both innate and adaptive immune protein networks that can be altered by DAC and shed light on the drug’s multifaceted immunomodulatory potentials to couple with various types of immunotherapy, including γδ T cells and NK cells, among others (*61–64*). We also demonstrate that DAC may enhance the polyfunctionality of the Vδ1 subtype to exert antitumor immunity. This finding differs from a previous report showing that DAC may upregulate the inhibitory receptor KIR2DL2/3 on *ex vivo* expanded Vδ2 T cells and inhibit their proliferation and effector functions (*65*). This disparity could be attributable to different γδ T cell subsets, expansion conditions, or DAC dosing schedules. As we gain additional insights into the phenotypic and functional heterogeneity of γδ T cells, caution should be taken when translating laboratory findings into the clinic.

In summary, our findings indicate the therapeutic potential of DAC in combination with adoptive immunotherapy with γδ T cells in lung cancer patients who are unresponsive to checkpoint inhibitors or lack targetable mutations/specific cancer antigens. In particular, those who have low ICAM-1-expressing tumors or unfavorable cytoskeleton gene signatures may benefit from epigenetic reprogramming to enhance the therapeutic efficacy of γδ T cells. This study also provides a molecular basis for pharmacologically modulating immune synapses to achieve effective antitumor immunity, which may facilitate the development of novel strategies in lung cancer management.

## MATERIALS AND METHODS

### Cancer cell lines and drug treatment

Human lung cancer cell lines, A549, H1299, HCC827, PC-9, PC-9-IR, H2981, H157, H1792, H2170, and colorectal cancer cell line, HCT116, were obtained from the American Type Culture Collection (ATCC). CL1-0 and CL1-5 human lung adenocarcinoma cell lines were kindly provided by Prof. Pan-Chyr Yang at the National Taiwan University College of Medicine (*66*). Cells were grown in HyClone PRMI medium (SH30027.02, GE Healthcare) supplemented with 10% GIBCO dialyzed FBS (26140-079, Life Technologies), 1% L-glutamine (A29168-01, Life Technologies), and 1% penicillin-streptomycin (15140-122, Life Technologies). Authentication of all cell lines used in this study was performed using short tandem repeat (STR) analysis. For drug treatment experiments, cells were cultured with 100 nM decitabine (DAC; A3656, Sigma-Aldrich) for three days with daily change of complete medium and drug replenishment, followed by 3 to 4 days of cell recovery in the decitabine-free medium. To block the DAC’s effect on immune synapse formation via actin cytoskeleton reorganization, the DAC-pretreated (D3R3) cells were treated with 1 μg/mL cytochalasin B (Cyto B; C6762, Sigma-Aldrich) for 1.5 hours prior to coculture with γδ T cells.

### SILAC labeling

For quantitative mass spectrometry analysis, lung cancer cells were cultured for at least seven cell doublings in SILAC RPMI medium (88365, Thermo Fisher Scientific) supplemented with 10% FBS and heavy isotope-labeled amino acids (L-Lysine 13C6, L-Arginine 13C6) (CLM-2247-H-1, CLM-2265-H-1, Cambridge Isotope Laboratories). DAC-treated cells grown in the regular medium were considered as SILAC light counterparts.

### Biotinylation and isolation of cell surface proteins

Cancer cells (around 1.5 x 10^7^ cells) were grown in a 15-cm culture dish and washed with ice-cold PBS (21040CM, Corning) before the labeling. A total of 5 mg of membrane-impermeable sulfosuccinimidyl-6-(biotin-amido) hexanoate reagent at the concentration of 0.3 mg/mL (EZ-link sulfo-NHS-SS-biotin; 21331, Thermo Fisher Scientific) was applied to the cells with gentle shaking at 4°C for 30 minutes. The reaction was quenched by 20 mM glycine (G8790, Sigma-Aldrich) in PBS for 10 minutes. Subsequently, biotinylated cells were collected by scraping in PBS containing 100 µM oxidized glutathione (G4376, Sigma-Aldrich) to prevent the reduction of the disulfide bridge in the labeling molecule. After centrifugation at 500 x g for 3 minutes, the cell pellet was subjected to a freezing-thawing step and resuspended in the lysis buffer [2% Nonidet P40 (NP40; 11332473001, Roche), 0.2% SDS (75746, Sigma-Aldrich), 100 µM oxidized glutathione and 1X Halt protease inhibitor cocktail (87786, Thermo Fisher Scientific) in PBS)] for 30 minutes on ice. The lysate then underwent sonication with five 15-second bursts and was centrifuged at 14,000 x g for 5 minutes at 4°C to remove insoluble materials. The protein concentration of the supernatants was determined using Pierce 660 nm Protein Assay Reagent Kit (22660, Thermo Fisher Scientific) supplemented with Ionic Detergent Compatibility Reagent (22663, Thermo Fisher Scientific). The biotinylated protein extracts from unlabeled and SILAC-heavy-labeled cancer cells were mixed in 1:1 (w/w), and the mixture was subjected to biotin-affinity purification. Pierce NeutrAvidin-agarose slurry (29200, Thermo Fisher Scientific) in 200 µl/mg of total proteins was conditioned by three washes in buffer A (1% NP40 and 0.1% SDS in PBS). Binding of biotinylated proteins was performed on a rotating mixer at 4°C overnight. The beads were then washed twice with buffer A, twice with buffer B [0.1% NP40, 10.3 M NaCl (4058-01, JT Baker) in PBS] and twice with buffer C (50 mM in PBS, pH 7.8). Proteins were eluted twice for 30 minutes each at 58°C with 150 mM dithiothreitol (DTT; D0632, Sigma-Aldrich), 1% SDS, 50 mM Tris-HCl in PBS (pH 7.8). Subsequently, 150 mM iodoacetamide (IAA; I6125, Sigma-Aldrich) was added to the sample followed by incubation for 30 minutes at room temperature in the dark to alkylate reduced cysteine residues in proteins. To concentrate the proteins and reduce the detergents in the membrane-rich eluate, we purified the eluate using an Amicon MWCO-10K centrifugation unit (UFC501096, Millipore) with the exchange buffer [0.5% SDS in 50 mM NH_4_HCO_3_ (09830, Sigma-Aldrich) aqueous solution]. The sample retained in the exchange buffer was then recovered for subsequent analyses (*67*).

### In-gel digestion and mass spectrometry

The purified biotinylated-surface proteins were fractionated using SDS-PAGE, followed by in-gel trypsin digestion, as previously described (*68*). The trypsinized peptides were vacuum-desiccated and solubilized in 0.1% trifluoroacetic acid (TFA; 299537, Sigma-Aldrich). The sample was then desalted using C18 Ziptip (ZTC18S960, Millipore) according to the manufacturer’s protocol. Peptide mass acquisition was performed on an Ultimate system 3000 nanoLC system connected to the Orbitrap Fusion Lumos mass spectrometer equipped with NanoSpray Flex ion source (Thermo Fisher Scientific). After loading the peptides into the HPLC instrument, the peptides were concentrated by a C18 Acclaim PepMap NanoLC reverse-phase trap column with a length of 25 cm and an internal diameter of 75 µm containing C18 particles sized at 2 µm with a 100 Å pore (164941, Thermo Fisher Scientific). The mobile phase aqueous solvent A (0.1% formic acid; 33015, Sigma-Aldrich) and organic solvent B [0.1% formic acid in acetonitrile (9829, JT Baker)] were mixed to generate a linear gradient of 2% to 40% solvent B for fractionated elution of peptides. The mass spectra were acquired from one full MS scan, followed by data-dependent MS/MS of the most intense ions in 3 seconds. Full MS scans were recorded with a resolution of 120,000 at m/z 200. For MS/MS spectra, selected parent ions within a 1.4 Da isolation window were fragmented by high-energy collision activated dissociation (HCD) with charge status of 2+ to 7+. The duration of the dynamic exclusion of parent ions was set at 180 seconds with a repeat count. Mass spectra were recorded by the Xcalibur tool version 4.1 (Thermo Fisher Scientific).

### LC-MS/MS data analysis and SILAC-based protein quantitation

The resulting Thermo RAW files were analyzed using MaxQuant v1.6.0.16 (*69*). MS/MS spectra were searched in the Andromeda peptide search engine against the UniProtKB/Swiss-Prot human proteome database as well as the reversed counterpart as a decoy database. The common contaminants list provided by MaxQuant software was applied during the search. Peptides with a minimum of 6 amino acids and maximum trypsin missed cleavages of 3 were considered. The setting of variable modifications includes methionine oxidation, protein N-terminal acetylation, asparagine/glutamine deamidation, and EZ link on primary amines after sulfhydryl reduction and alkylation by IAA (EZ-IAA, +145.020 Da) (*70*). Carbamidomethyl cysteine was applied as a fixed modification. Initial parent peptide and fragment mass tolerance were set to 20 ppm and 0.5 Da, respectively. False discovery rate (FDR) filtration of the peptide-spectrum match and protein assignment were utilized at 0.05 and 0.01, respectively. Finally, proteins identified as a reverse decoy, matched with only one unique peptide, and as common contaminations were excluded before further analysis. The Thermo RAW files and MaxQuant results have been deposited to the ProteomeXchange Consortium with the dataset identifier MSV000084997 through the MassIVE partner repository.

### Patient-derived lung cancer cells from malignant pleural effusions (MPE**s)**

The MPEs from lung cancer patients were centrifuged at 1,800 x g for 5 minutes to collect cell pellets. Cell pellets were resuspended in 4 ml PBS and subject to density gradient centrifugation with Ficoll-Paque PLUS (17144002, GE Healthcare) according to the manufacturer’s instructions. Briefly, the resuspended cells were carefully loaded onto 3 ml Ficoll-Paque PLUS in a 15-ml centrifuge tube and layered by centrifugation at 1,800 x g for 20 minutes at room temperature. Nucleated cells enriched at the interface were collected and washed by at least three volumes of PBS. Then the cells were pelleted by centrifugation at 300 x g for 5 minutes. Red blood cells (RBC) in the cell pellet were lysed with RBC lysis buffer [155 mM NH_4_Cl (11209, Sigma-Aldrich), 10 mM KHCO_3_ (2940-01, JT Baker) and 0.1 mM EDTA (34550, Honeywell Fluka) in deionized water], and discarded after centrifugation of the sample at 300 x g for 3 minutes. The cell pellet was finally washed twice with PBS and centrifuged at 300 x g for 3 minutes. The collected cells were grown in DMEM/F-12 (1:1 in v/v; 11330, Thermo Fisher Scientific) supplemented with 5% FBS, 2% penicillin-streptomycin, 0.4 μg/mL hydrocortisone (H088, Sigma-Aldrich), 5 μg/mL insulin (I2643, Sigma-Aldrich), 10 ng/mL epidermal growth factor (PHG0311L, Invitrogen), 24 μg/mL adenine (A2786, Sigma-Aldrich) and 6 μM Y-27632 (ALX-270-333, Enzo Life Sciences). Floating lymphocytes in the primary culture were eliminated by PBS washing before a detachment of adherent cells with trypsin-EDTA (25200072, Thermo Fisher Scientific) at 0.25% in PBS during each passage. The removal of fibroblasts was achieved due to their faster adhesion to the culture dish than tumor cells. We transferred the trypsinized-cell suspension to a new culture dish and let it sit at 37°C until the two types of cells were separated from each other. After repeated subcultures, the purity of the tumor cell population was confirmed by measuring the surface expression of EpCAM by flow cytometric analysis (*71*).

### Isolation and *ex vivo* expansion of γδ T lymphocytes

1×10^7^ Peripheral blood mononuclear cells (PBMC) obtained from healthy donors were seeded in each well coated with anti-TCR PAN γ/δ antibody (IMMU510 clone) using a 6-well culture plate. The culture medium contained Optimizer™ CTS™ T-Cell Expansion SFM (A1048501, Thermo Fisher), 15 ng/mL IL-1β (AF-200-01B), 100 ng/mL IL-4 (AF-200-04), 7 ng/mL IL-21 (AF-200-21), and 70 ng/mL IFN-γ (AF-300-02, Peprotech). After seven days, the medium was changed to Optimizer CTS medium, 5% human platelet lysate (PLS1, Compass Biomedical), 70 ng/mL IL-15 (AF-200-15, Peprotech), and 30 ng/mL IFN-γ. These cells were harvested on Day 21 for subsequent experiments, including *in vitro* cytotoxicity and animal experiments. Informed consent was obtained from individual healthy donors before enrollment. The study was approved by the Institutional Review Board (IRB) of National Taiwan University Hospital.

### Immunophenotypic and functional analysis of γδ T cells

*Ex vivo* expanded γδ T cells were stained with immunofluorescence antibodies targeting the surface markers, including γδTCR Vδ1, γδTCR Vδ2, CD27, CD69, NKG2D, TGF-β1, and CD107a (**table S1**). Subsequently, γδ T cells were fixed and permeabilized using Cytofix/Cytoperm solution (554714, BD Biosciences) for 20 minutes at 4°C for intracellular staining of cytokines, including IL-2, IL-10, IL-17A, IFN-γ, and TNF-α (**table S1**). Staining was performed at 4°C for 30 minutes in the dark. The samples were washed and fixed with 100 μl of 1X IOTest3 Fixative Solution (A07800, BECKMAN COULTER) per well for at least 10 minutes at 4°C. Cells were then resuspended in 300 µl PBS and analyzed using a BD LSR Fortessa flow cytometry (BD Biosciences). Acquired data were analyzed using FlowJo software (Tree Star). For polyfunctional response measurement, γδ T cells were stimulated with 30 ng/mL phorbol 12-myristate 13-acetate (PMA; P1585, Sigma-Aldrich) and 1 μg/mL ionomycin (I9657, Sigma-Aldrich) in the presence of monensin (00-4505-51, eBioscience) and brefeldin A (420601, BioLegend) for 4 hours at 37°C. After stimulation, γδ T cells were transferred to v-bottom 96-well plates and stained for surface markers and intracellular cytokines as described above. As an unactivated control, γδ T cells were incubated only with dimethyl sulfoxide (DMSO; D2650, Sigma-Aldrich), monensin, and brefeldin A before staining.

### Single-cell mass cytometry (CyTOF)

Samples were processed as described with few modifications (*72*). Briefly, the cell samples were first stained for viability with cisplatin (201064, Fluidigm) and then fixed with 1.5% paraformaldehyde (15710, Electron Microscopy Sciences) at room temperature for 10 minutes followed by two washes with Cell staining medium (CSM) [PBS containing 0.5% bovine serum albumin (BSA; A3059, Sigma-Aldrich) and 0.02% sodium azide (S2002, Sigma-Aldrich)]. Formaldehyde-fixed cell samples were then subjected to pre-permeabilization palladium barcoding, as previously described (*73*). The barcoded samples were first incubated with anti-TCR Vδ 1-FITC for 30 minutes on ice, washed once with CSM, and then stained with metal-conjugated antibodies against surface markers for 1 hour. After incubation, samples were washed once with CSM, permeabilized with 1x eBioscience™ Permeabilization Buffer (00-8333-56, Thermo Fisher Scientific) on ice for 10 minutes, and then incubated with metal-conjugated antibodies against intracellular molecules for 1hour. Cells were washed once with 1x eBioscience™ Permeabilization Buffer and then incubated at room temperature for 20 minutes with an iridium-containing DNA intercalator (201192A, Fluidigm) in PBS containing 1.5% paraformaldehyde. After intercalation/fixation, the cell samples were washed once with CSM and twice with water before measurement on a CyTOF mass cytometer (Fluidigm). Normalization for detector sensitivity was performed as previously described (*74*). After measurement and normalization, the individual files were debarcoded (*73*) and gated according to **Figure S3A**. viSNE maps were generated using software tools available at https://www.cytobank.org/. For antibody conjugations, antibodies in carrier-free PBS were conjugated to metal-chelated polymers (**table S1**) according to the manufacturer’s protocol. Antibody conjugation to bismuth was carried out as previously described (*75*). The CyTOF raw FCS files have been deposited to FlowRepository database with the identifier FR-FCM-Z2G5.

### Data analysis for mass cytometry data

For the comparison between *ex vivo* expanded γδ T cells with and without DAC treatment, 50,000 cells from each group were randomly sampled and pooled together for clustering using X-shift (*76*), a density-based clustering method. All markers except the ones used for gating were selected for clustering. Clusters separated by a Mahalanobis distance less than 2.0 were merged. The optimal nearest-neighbor parameter, K, was determined as 20 using the elbow method. The expression level and the cell frequency in each cluster were exported and represented by heatmaps and piecharts using R. For the correlation between expression levels of markers in CD3+ T cells at baseline and after *ex vivo* expansion, pairwise Pearson correlation coefficients were calculated. The heatmap was generated and clustered based on hierarchical clustering of the Pearson correlation coefficients by R.

### γδ T cell-mediated cytotoxicity assays

Cancer cells were cocultured with γδ T cells at an effector to target (E:T) ratio of 3:1 at 37°C for 2 hours. After coculture, cell death was evaluated by flow cytometric analysis using FITC Annexin V Apoptosis Detection Kit I (556547, BD Biosciences). In addition, we performed real-time monitoring of γδ T-mediated killing of cancer cells using the Electric Cell-Substrate Impedance Sensing (ECIS) monitoring system with an 8W10E+ culture chamber (Applied Biophysics). Mock or DAC-treated cancer cells were cultured in the chamber at 37°C overnight until the cancer cells were fully attached to the bottom of the wells. Following the addition of γδ T cells, the detachment of cancer cells indicating cell death was recorded in real-time using multiple frequency capture with the ECIS software. Relative impendence at the time of γδ T cell addition was used for between-sample normalization. For non-contact killing experiments, cancer cells were seeded in the bottom wells of a 24-well Transwell system overnight, followed by the addition of γδ T cells onto the top 0.4 µm pore membrane inserts that are impermeable to cells (353095, Falcon). The coculture was performed at an E:T ratio of 10:1 at 37°C for overnight. The death of cancer cells in the bottom wells was evaluated by flow cytometric analysis using FITC Annexin V Apoptosis Detection Kit I.

### γδ T cell chemotaxis assay

Cancer cells were cocultured with γδ T cells in a Transwell system with a 3 µm pore membrane insert (3415, Falcon) that allows γδ T cells to pass through. Prior to the coculture, cancer and γδ T cells were stained with vital dyes, Calcein AM (1755, BioVision) and Hoechst 33342 (H3570, Thermo Fisher Scientific), respectively. After coculture for 2 hours, the γδ T cells (positive for Hoechst 33342) that had migrated into the bottom chamber were imaged under a fluorescence microscope and quantified by ImageJ.

### Immunofluorescence imaging of immune synapses

Cancer and γδ T cells were spun down onto poly-L-lysine coated coverslips by using a Cyto-Tek table-top cytofuge at 500 rpm for 5 minutes (Sakura Scientific) before fixation with ice-cold methanol or 4% (w/v) paraformaldehyde in PBS for 10 minutes. The fixed samples were blocked in detergent-free blocking solution [1% normal donkey serum (ab7475, abcam) and 3% BSA in PBS] without membrane permeabilization by detergents. Subsequently, cells were incubated with primary antibodies diluted in the blocking solution for 1 hour in a moist chamber at room temperature and washed with PBS. Labeling of fluorescent secondary antibodies at 1:200 dilution in PBS was carried out in the blocking buffer for 1 hour at room temperature. After PBS washing, cell nuclei were stained with Hoechst 33342 or DAPI (62248, Thermo Fisher Scientific) in PBS. Finally, the samples were washed with PBS and mounted on the slides. All slides were examined under an epifluorescence microscope EVOS FLc (Invitrogen), and the images were analyzed using the ImageJ software. For the visualization of immune synaptic proteins, cells were imaged with a high-resolution confocal microscope (LSM780, Zeiss) with a 63X oil objective and analyzed with the confocal software ZEN (Zeiss). Antibodies used in the study are listed in **table S1**.

### Overexpression and CRISPR/Cas9 gene knockout experiments

In overexpression experiments, doxycycline-inducible ICAM-1 overexpressing vector was created through the cloning of full-length *ICAM1* cDNA into a Tet-On lentiviral plasmid pLVX-Tight-Puro (632162, Clontech). The gene expressing vector and the regulator vector (pLVX-Tet-On Advanced) were packaged with VSV-G pseudotyped lentivirus particles in 293 T cells. After co-transfection of the two lentiviral particles into lung cancer cells, the cells were then grown in the selection media containing G418 (10131035, Thermo Fisher Scientific) and puromycin (A1113803, Thermo Fisher Scientific) at proper concentrations for the retention of both plasmids. In knockout experiments, editing of the *ICAM1* genome locus on H1299 cells was achieved through coexpression of the Cas9 protein with the guide RNAs (gRNAs) targeting to *ICAM1* exon 2 at the sequences 5’-TCAAAAGTCATCCTGCCCCG-3’ and 5’-GTGACCAGCCCAAGTTGTTG-3’. ICAM-1-null cell lines were established through clonal propagation from single cells. The overexpression and loss of ICAM-1 protein were validated by flow cytometry with an anti-ICAM1 antibody (BBA20, R&D Systems). Besides, the edited genome patterns at *ICAM1* locus around the gRNA-targeting sites of the ICAM1-knockout lines were validated by Sanger sequencing after PCR amplification (see **table S1** for the PCR primer pair). Analysis of the sequencing results for the CRISPR/Cas9-edited *ICAM1* genome locus is performed using multiple sequence alignment tool (ClustalO) on QIAGEN CLC Genomics Workbench software (v20.0.2).

### RNA preparation and mRNA-seq analysis

Total RNA was extracted with the PureLink RNA Mini Kit according to the manufacturer’s instructions (12183018A, Invitrogen). The quality of RNA was evaluated using a Bioanalyzer 2100 with RNA 6000 Nano LabChip kit (5067-1511, Agilent Technologies). mRNA-seq libraries were prepared using TruSeq Stranded mRNA Library Prep Kit (RS-122-2101, Illumina) and sequenced using the HiSeq 4000 system. Raw reads were processed with adapter trimming and quality filtering using *Trimmomatic* with default settings (*77*). The cleaned reads were aligned to UCSC human genome hg19 using *RSEM* tool with the *bowtie2* aligner (*78, 79*). Mapped reads were counted for each gene using the R packages GenomicFeatures (v1.36.4) and GenomicAlignments (v1.20.1), according to the GENCODE human GRCh37 annotation (https://www.gencodegenes.org/human/release_25lift37.html) (*80*). Finally, FPKM normalization of the raw-counts was performed using DESeq2 (v1.24.0) (*81*). Bar graphs and dot plots of RNA-seq data were created by ggplot2 (v3.2.1). Networks and upstream regulator analysis were conducted by Ingenuity Pathway Analysis (IPA^®^; version 01-16, Qiagen). The BAM files with alignments and analysis results of mRNA-seq data have been deposited to Gene Expression Omnibus (GEO) database with the identifier GSE145663.

### Gene ontology and gene set enrichment analysis

Surface proteins upregulated by more than 1.4-fold with DAC treatment in the quantitative membrane proteomic analysis were subjected to gene ontology (GO) analysis using the AmiGO 2 web tool, PANTHER (v2.5.12), with Fisher’s Exact test. For gene set enrichment analysis (GSEA), we used mRNA-seq data of cancer cell lines with and without DAC treatment to identify gene sets enriched in the DAC-treated samples (FDR < 0.25 and nominal p-value < 0.05). Cytoskeleton-associated gene sets are retrieved from the Molecular Signatures Database (MSigDB, v6.2).

### Genome-wide DNA methylation analysis

Genomic DNA of cancer cells was extracted with the QIAamp DNA Mini Kit (51304, QIAGEN) according to the manufacturer’s instruction. The DNA concentration and quality were evaluated by NanoDrop 2000 (Thermo) and electrophoresis with 0.8% agarose gel, respectively. Bisulfite conversion of 1 µg genomic DNA was performed using EZ DNA Methylation Kit (D5001, Zymo Research). The bisulfite-converted DNA samples were subject to genome-wide methylation analysis using the Illumina Infinium MethylationEPIC BeadChips. Raw intensity data were obtained as IDAT files and processed using R package *minfi* v1.30.0 (*82*) with a probe annotation package for Illumina EPIC array (IlluminaHumanMethylationEPICanno.ilm10b4.hg19). The data were quantile normalized using *the preprocessQuantile* function of *minfi*. We removed low-quality probes with detection P-value > 0.01 as well as probes described as single nucleotide polymorphisms (SNPs), cross-reactive and genetic variants (*83, 84*). Finally, a total of 692,476 probes were used for further analysis. The signal intensity raw IDAT files and analysis results of MethylationEPIC data have been deposited to GEO database with the identifier GSE145588.

### Genome-wide chromatin accessibility analysis

Chromatin accessibility of lung cancer cells before and after DAC treatment was analyzed using the Omni-ATAC protocol (*44*) with modifications. After harvesting cells with trypsin/EDTA, a total of 1×10^5^ cells were resuspended in 1 ml of cold ATAC resuspension buffer [ATAC-RSB; 10 mM Tris-HCl pH 7.4, 10 mM NaCl, and 3 mM MgCl_2_ (AM9530G, Thermo Fisher Scientific) in water] supplemented with 0.1% Tween-20 (P2287, Sigma-Aldrich). Cells were centrifuged at 650 x g for 5 minutes at 4°C to remove the buffer and lysed in 50 μl of ATAC-RSB containing 0.1% NP40, 0.1% Tween-20, and 0.01% digitonin (G9441, Promega) on ice for 3 minutes. Subsequently, the lysate was washed by 1 ml of ATAC-RSB containing 0.1% Tween-20, and the supernatant was discarded following centrifugation at 650 x g for 10 minutes at 4°C to pellet the cell nuclei. The nuclear fraction was then resuspended in 50 µl of 1:1 (v/v) premix of 2X Tagmentation DNA (TD) buffer [20 mM Tris-HCl, 10 mM MgCl_2_, 10% dimethyl formamide (D4551, Sigma-Aldrich), pH 7.6] and Transposition buffer [2 µl of Illumina adapters-bearing Tn5 transposase (FC-121-1030, Illumina), 0.02% digitonin and 0.2% Tween-20 in PBS]. The sample was incubated at 37°C for 30 minutes in a water bath and vortexed every 5 minutes to facilitate the transposition reaction. At the end of the reaction, the transposed DNA fragments were harvested and cleaned up with Zymo DNA Clean and Concentrator-5 kit (D4011, Zymo Research). The libraries were indexed and amplified with Illumina i5/i7 indexing primers (FC-121-1011, Illumina) by PCR reaction to reach a target concentration of 4 nM in 20 μl. Library quality was checked by Agilent 2100 Bioanalyzer with high sensitivity DNA kit (5067-4626, Agilent Technologies). The sample was sequenced on the Illumina HiSeq X Ten platform with 151 bp paired-end sequencing for an average of 60 million raw reads per sample.

### Bioinformatic analysis of the Omni-ATAC-seq data

The Nextera adapter sequence (5’-CTGTCTCTTATACACATCT-3’) was trimmed from the raw reads using CutAdapt v2.7 (*85*). Trimmed reads of each sample were mapped to the human reference genome build UCSC hg19 using BWA mem v0.7.17-r1188 (*86*) with default -M parameter and a maximum fragment length of 2,000. The PCR duplicated reads were marked using Picard (v2.21.4, http://broadinstitute.github.io/picard/). Further quality filtering was performed using SAMtools (v1.9, PMC2723002) to remove unmapped reads, unmapped mates, PCR duplicated reads, unpaired alignments, and reads mapped to mitochondrial DNA. The size distribution of the filtered reads and nucleosome-occupancy frequency was evaluated by ATACseqQC v1.8.5 (*87*). The broad peak regions of ATAC-seq were called using MACS2 v2.2.5 (*45*) with parameters: --nomodel --shift -75 --extsize 150 --keep-dup all --broad --broad-cutoff 0.1. Annotation of the peaks and the distance of each peak to the closest TSS were determined using ChIPseeker v1.20.0 (*88*). The bam files of the post-filtering read alignment from mock- and DAC-treated samples were simultaneously normalized with reads per genomic content (RPGC) approach by --effectiveGenomeSize 2827437033 and the log2 ratio of decitabine/mock per bin was reported using deepTools v3.3.1 (*89*). The BAM files with genome mapping and MACS2-called broad peaks results of ATAC-seq data have been deposited to GEO with the identifier GSE145663.

### Transcriptomic data of primary lung cancer tissues

The genome-wide gene expression data by mRNA-seq were obtained from two lung adenocarcinoma cohorts, National Taiwan University Hospital (NTUH), and the Cancer Genome Atlas (TCGA) database. The NTUH data were generated by our laboratory and deposited in the Gene Expression Omnibus (GEO) database with an accession number of GSE120622. The curated TCGA lung adenocarcinoma (LUAD) data of normalized gene expression (RNASeq2GeneNorm) and clinical information were acquired using an R package, curatedTCGAData v1.6.0 (*90*). Heatmaps of the Z-transformed gene expression level of selected genes of NTUH and TCGA mRNA-seq data were created using the R package pheatmap (v1.0.12).

### Combination therapy of DAC and γδ T cells in a tumor xenograft mouse model

Six-week-old male NOD.*Cg-PrkdcscidIl2rgtm1Wjl*/SzJ (NSG) mice were purchased from the National Laboratory Animal Center (Taiwan) and maintained under the standard pathogen-free condition. H1299 lung cancer cells were injected subcutaneously into the mice (1×10^7^/mouse). The therapy was started at seven days post-injection. For each two-week treatment cycle, tumor-bearing mice were treated with DAC (0.2 mg/kg BW) by intraperitoneal injection on Day 1, 2, and 3. Human γδ T cells (1×10^7^/mouse) were intravenously injected via tail vein on Day 5 and 10. The treatment was continued until the death of mice or the end of experiments. The survival time of individual mice in each treatment group was recorded. The surviving mice on day 42 post-tumor injection were sacrificed to obtain tumors for pathologic examination and hematoxylin and eosin (H&E) staining. All mice experiments were approved by the NTU College of Medicine Institutional Animal Care and Use Committee (IACUC) (Protocol #20180077).

### Statistical analysis

Statistical analysis was performed using GraphPad Prism 8 and computing environment R. Mann-Whitney or unpaired t-tests were used to compare the means between two groups, whereas one-way ANOVA with Tukey’s multiple comparisons was used to compare the means of three or more groups. Overall survival analysis for the NTUH and TCGA lung adenocarcinoma cohorts was performed using the Cox regression model, as well as the Kaplan–Meier method. Calculation and plotting of the survival curve were performed using R packages survival (v3.1-8) and survminer (v0.4.6), respectively. Heatmaps of the Z-transformed gene expression level of mRNA-seq data were created by using R package pheatmap (v1.0.12). Bar and spot charts representing methylation ß-value and RNA-seq FPKM were created by ggplot2 (v3.2.1).

## ACKNOWLEDGMENTS

We sincerely thank Dr. Stephen Baylin at Johns Hopkins University School of Medicine for his valuable feedback on the manuscript. We acknowledge the service provided by the Flow Cytometric Analyzing and Sorting Core and the Imaging Core of the First Core Laboratory and the Laboratory Animal Center at National Taiwan University College of Medicine. We would also like to thank the Tai-Cheng Cell Therapy Center and the Instrumentation Center of National Taiwan University for technical assistance. Mass cytometry analyses were performed by GRC Mass Core Facility of Genomics Research Center, Academia Sinica, Taipei, Taiwan, and Stanford Human Immune Monitoring Center, USA. This study was supported by Ministry of Science and Technology (MOST 105-2628-B-002 -040), National Health Research Institutes (NHRI-EX106-10610BC), Excellent Translational Medical Research Grant by National Taiwan University College of Medicine and National Taiwan University Hospital (105C101-71, 106C101-B1), as well as Institutional Top-down Research Grant by National Taiwan University Hospital (NTUH 107-T10, 108-T10, 109-T10).

---

**Supplementary Figure 1.**
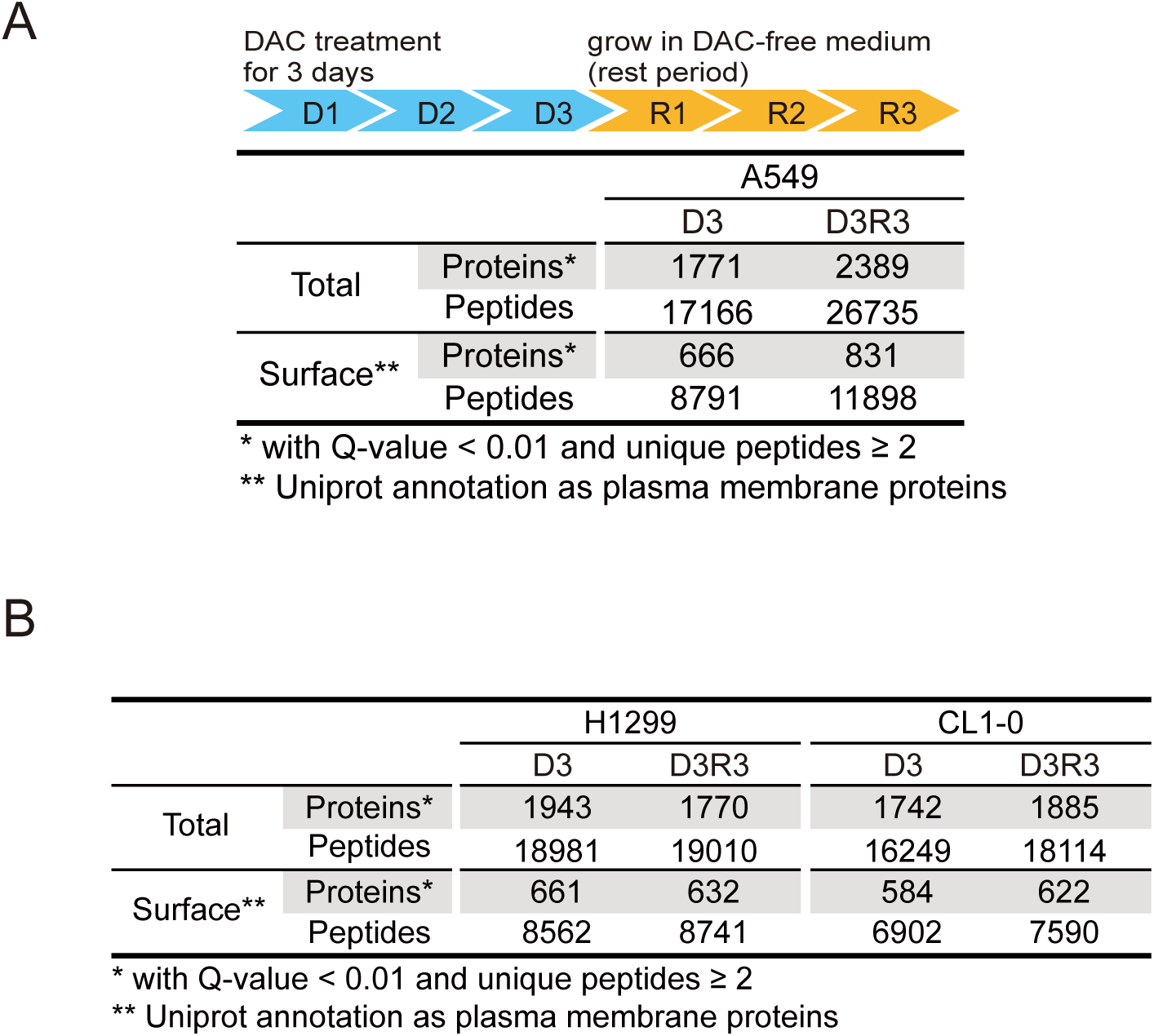
Summary of the identified plasma membrane proteomes. (**A**) Numbers of identified total and surface proteins/peptides in A549 human lung cancer cells subject to 100 nM decitabine daily treatment for 72 hours (D3), followed by growing in drug-free medium for three days (D3R3). (**B**) Numbers of identified total and surface proteins/peptides in H1299 and CL1-0 human lung cancer cells subject to 100 nM decitabine daily treatment for 72 hours (D3), followed by growing in drug-free medium for three days (D3R3).

**Supplementary Figure 2.**
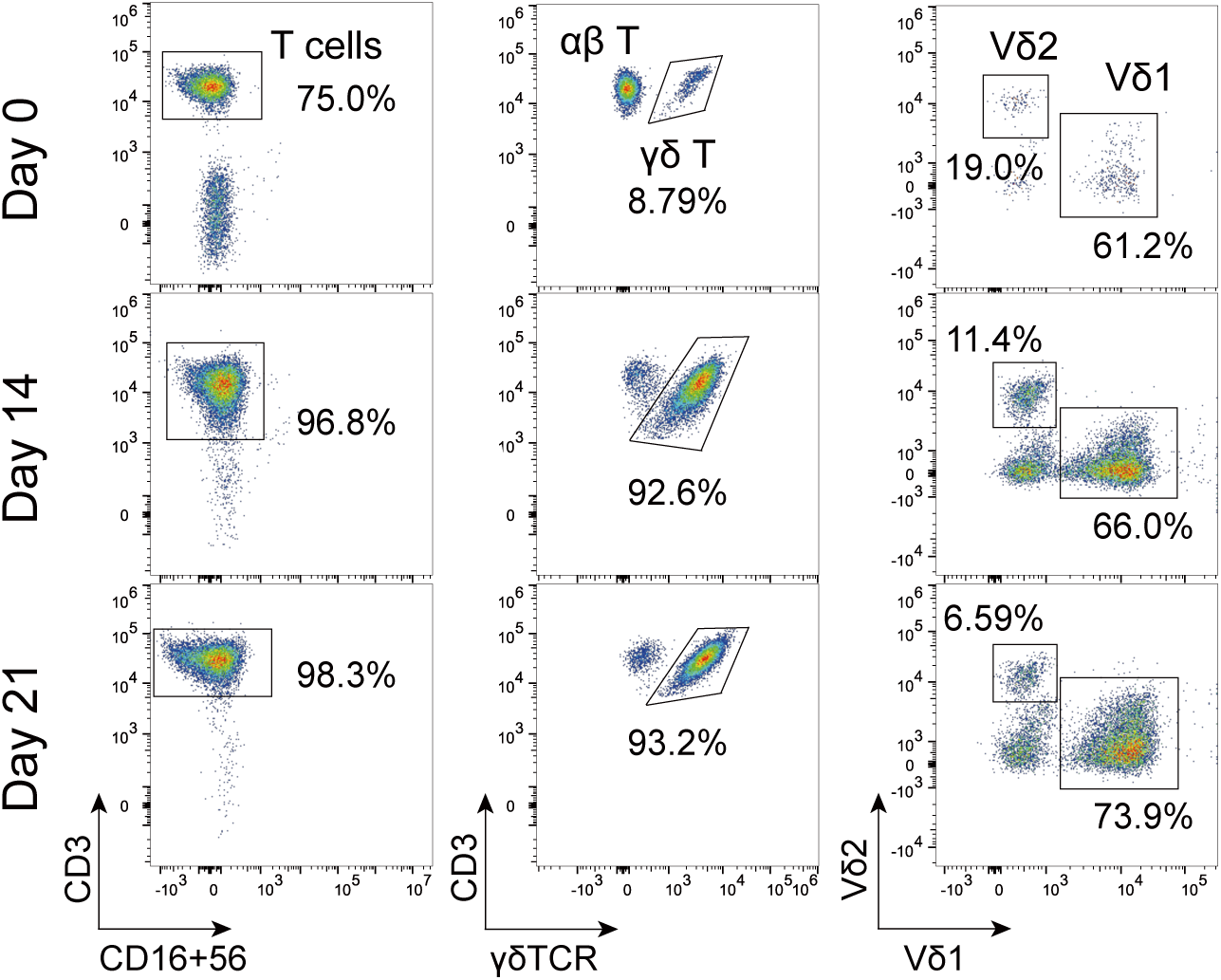
Gating strategy used to assess the *ex vivo* expanded γδ T lymphocytes. A representative plot of flow cytometric analysis showing *ex vivo* expansion of γδ T cells from peripheral blood mononuclear cells of a healthy donor at day 0, 14, and 21.

**Supplementary Figure 3.**
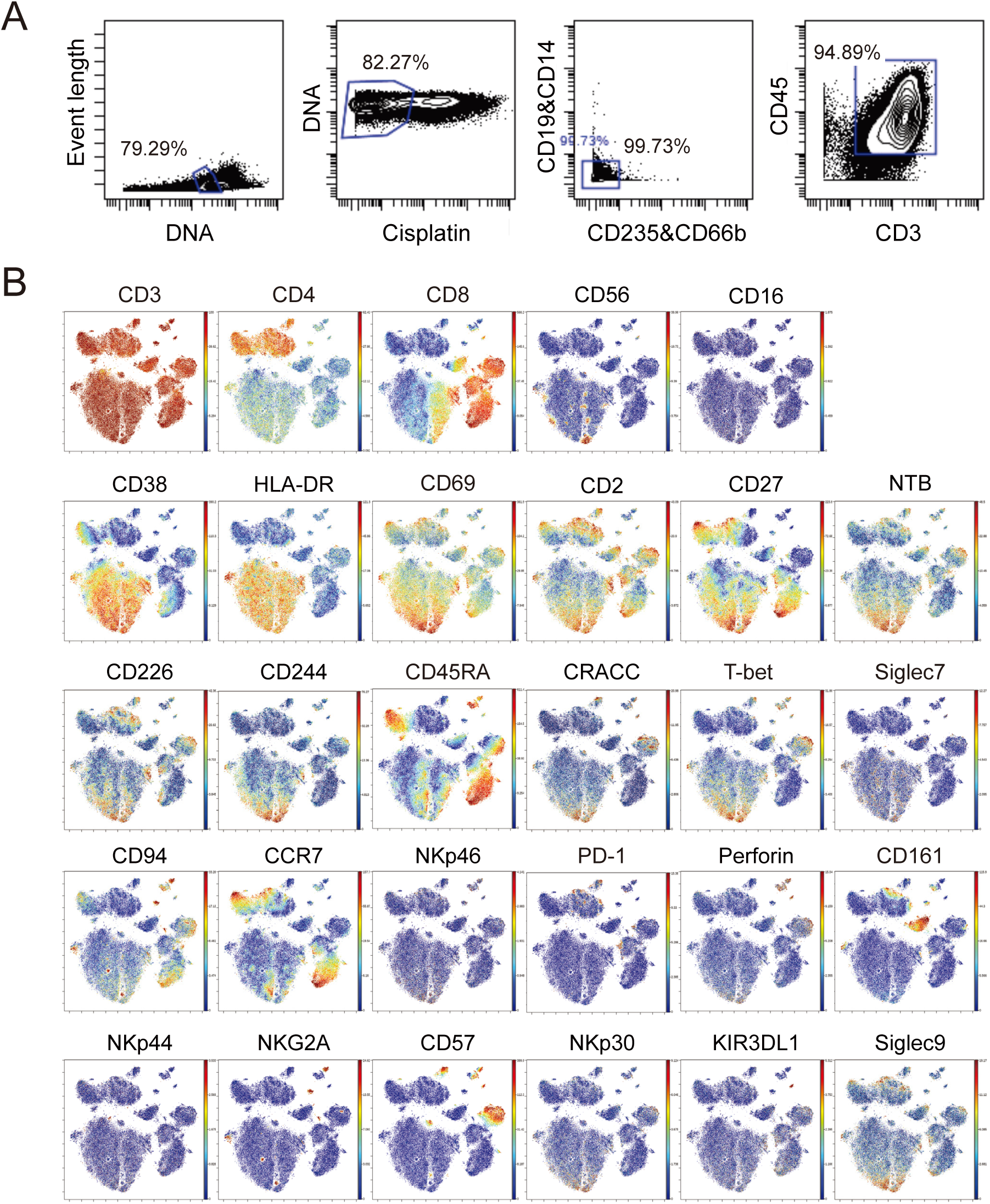
Single-cell mass cytometry analysis of CD3^+^ T cells before and after the expansion of γδ T cells. (**A**) A representative gating strategy to identify CD3+ T cells. Doublet and cell debris are excluded based the DNA content. Cell viability is determined by cisplatin stain. B cells, monocytes, granulocytes and red blood cells are excluded using CD19 and CD14, CD66b, and CD235, respectively. (**B**) An extended panel of tSNE plots for CD3+ T cells from PBMC at baseline and after *ex vivo* γδ T cell expansion analyzed by mass cytometry (CyTOF). Each dot represents a single cell. For individual markers, the color represents the expression level of the indicated markers. Red is high and blue is low.

**Supplementary Figure 4.**
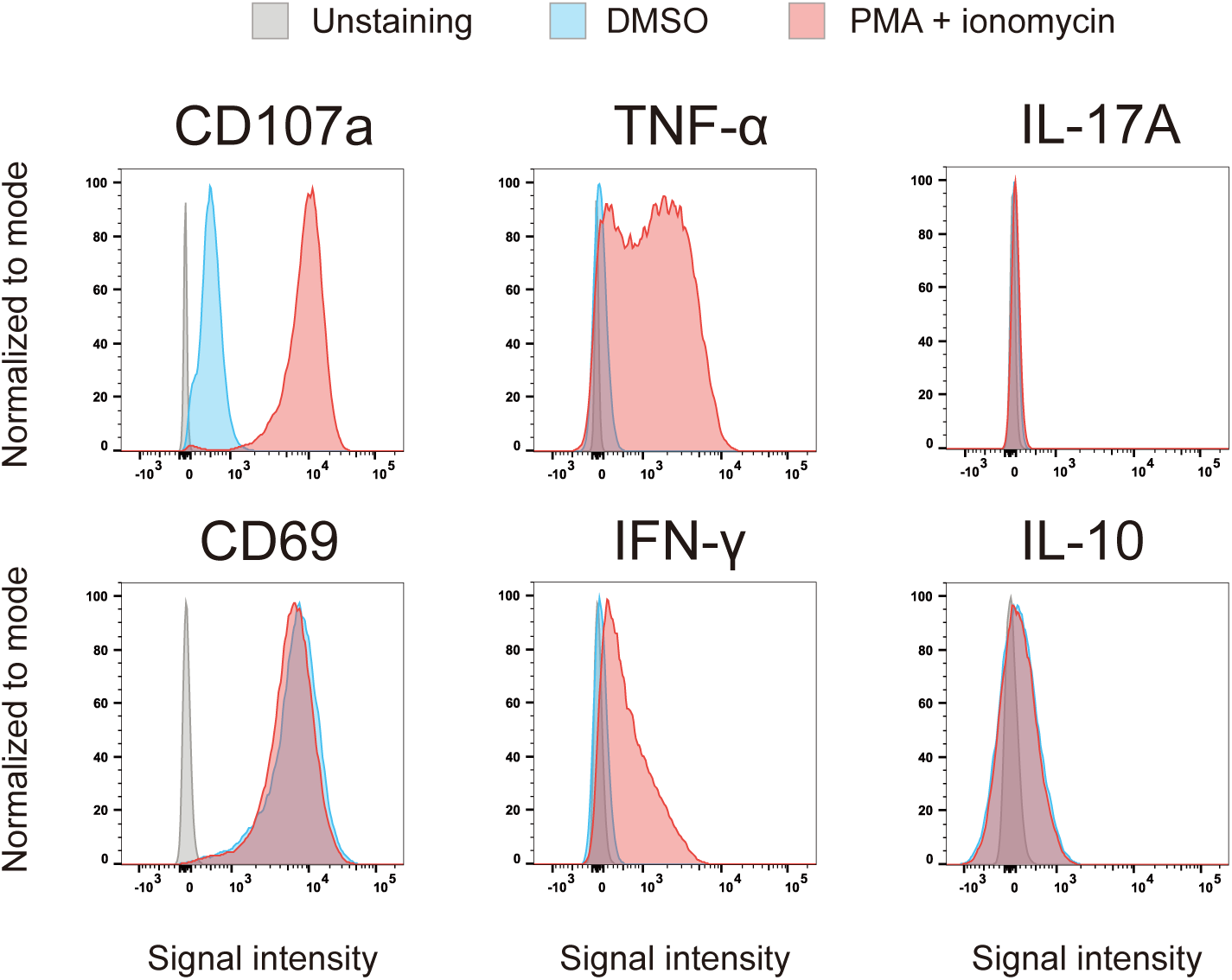
Validation of the markers characterized by CyTOF. Flow cytometric analysis of T cell activation markers (CD69, CD107a), antitumor effector cytokines (TNF-α, IFN-γ) and protumor effector cytokines (IL-17A, IL-10) produced by Vδ1 cells following PMA (30 ng/ml) and ionomycin (1 μg/ml) stimulation for 4 hours.

**Supplementary Figure 5.**
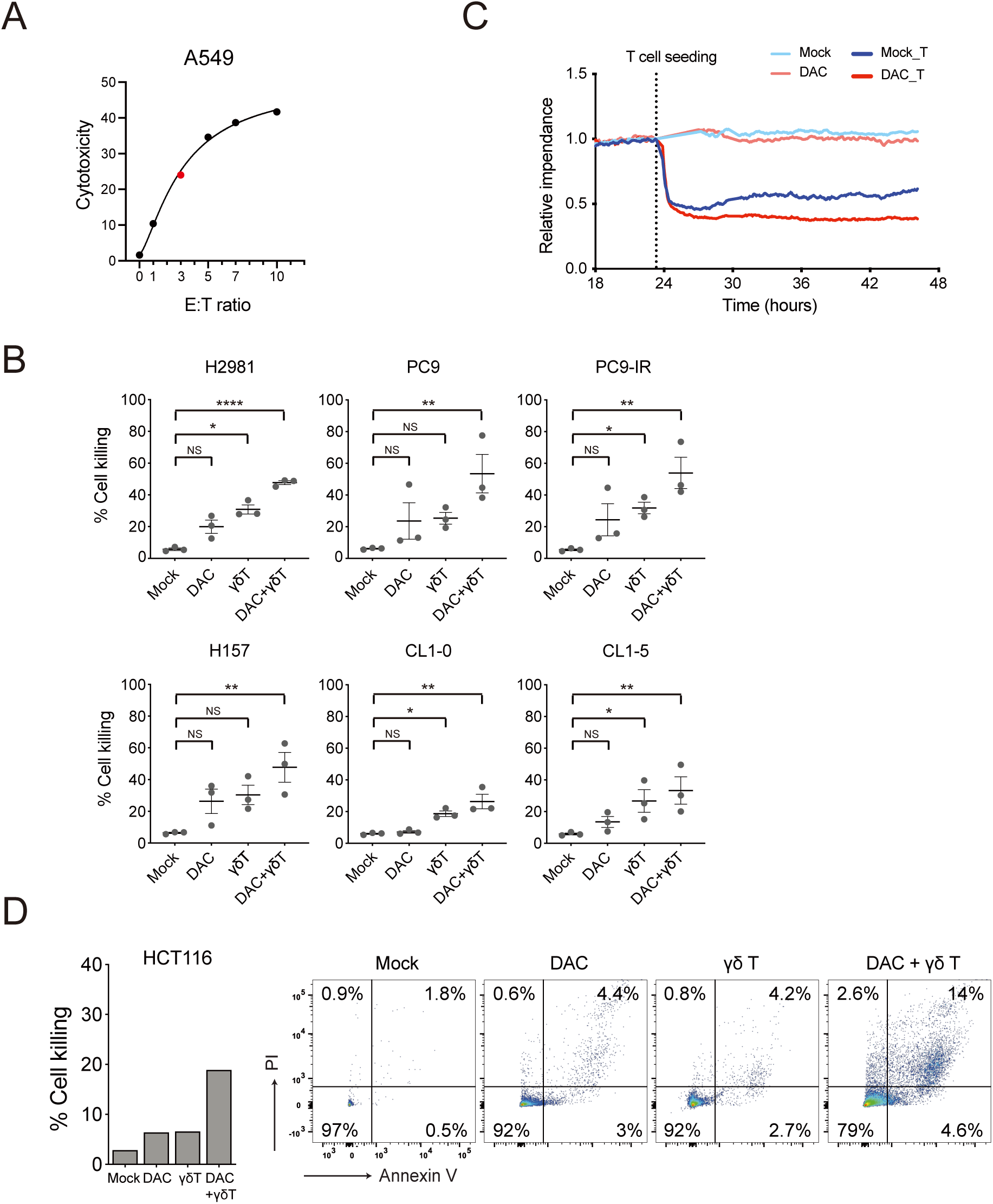
Decitabine-synergized cytotoxicity of cancer cells by γδ T. (**A**) Cytolysis of A549 lung cancer cells by γδ T cells at effector to target (E:T) ratio of 1, 3, 5, 7, and 10 is determined by calcein-release assay. (**B**) Annexin V and propidium iodide apoptosis assays of human lung cancer cell lines — H2981, PC9, PC9-IR, H157, CL1-0, and CL1-5 — upon treatments with 100 nM DAC alone, γδ T cells alone or DAC/γδ T cells combination. E:T ratio is 3:1. Data represent three biological replicates presented as mean ± SEM. *p* value is calculated by one-way ANOVA with Tukey’s multiple comparison test (*, *p* < 0.05; **, *p* < 0.01). (**C**) Real-time impedance-based cell viability measurement of A549 lung cancer cells subject to mock, DAC alone, γδ T alone or combination of DAC and γδ T treatment using the electric cell-substrate impedance sensing (ECIS^TM^) system. Acquired raw data are normalized by the impedance at the latest time prior to adding γδ T cells of each condition. DAC treatment alone has minimal effects on cell viability but may potentiate γδ T-mediated cytolysis when combined with γδ T. (**D**) Bar graphs showing Annexin V and propidium iodide (PI) apoptosis assays of HCT116 colorectal cancer cells upon treatments with 100 nM DAC alone, γδ T cells alone or DAC/γδ T cells combination. Representative flow cytometric analysis is shown on the right.

**Supplementary Figure 6.**
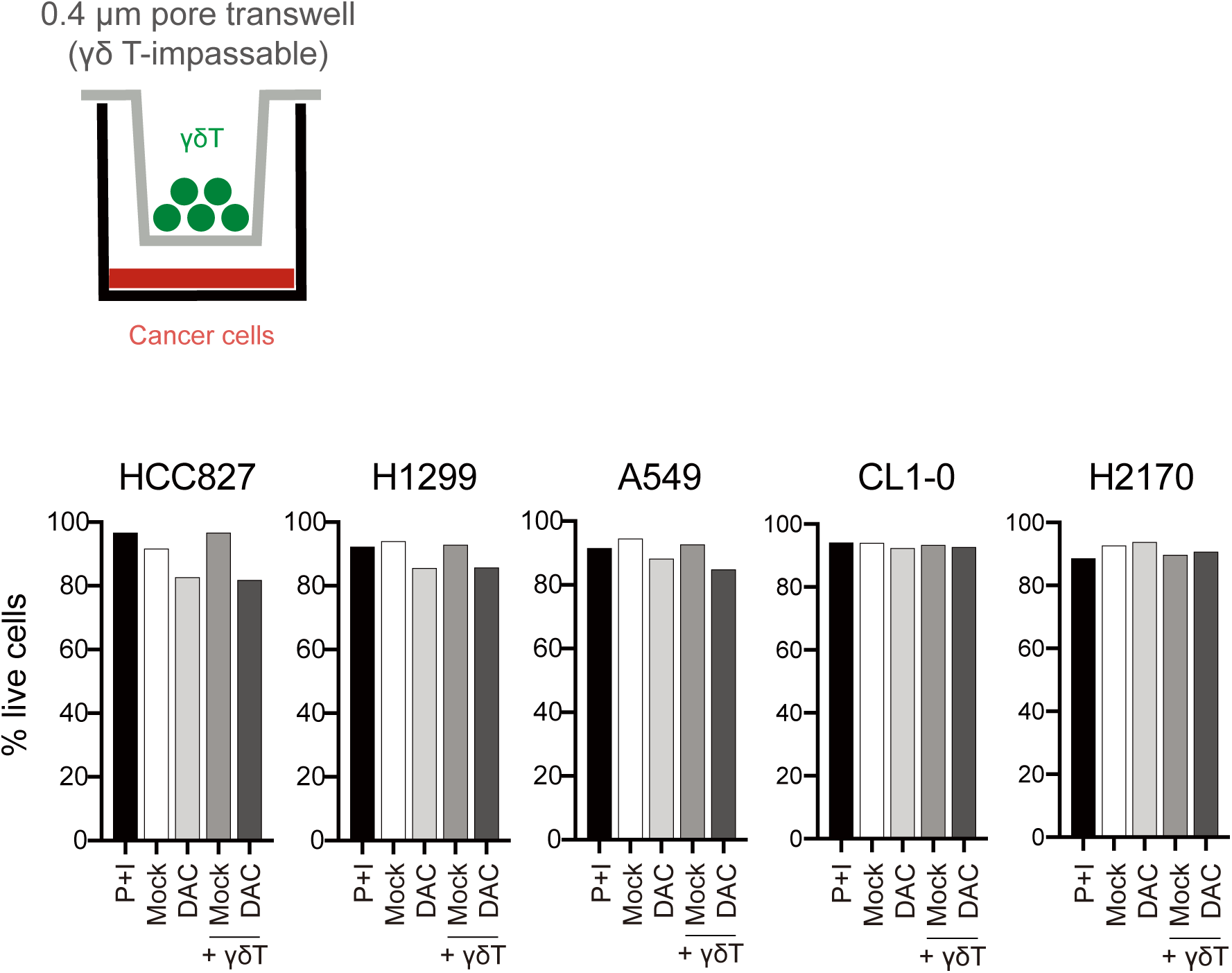
Decitabine does not trigger non-contact cytotoxicity of cancer cells by the *ex vivo* expanded γδ T cells. Lung cancer cell lines are pretreated with DAC for 72 hours and rest for 3 days before coculture with γδ T cells in a Transwell coculture system permeable to cytokines and cytotoxic mediators (lower chamber: lung cancer cells; upper chamber: γδ T cells). Cell death is analyzed by Annexin V apoptosis assays after coculture for 24 hours.

**Supplementary Figure 7.**
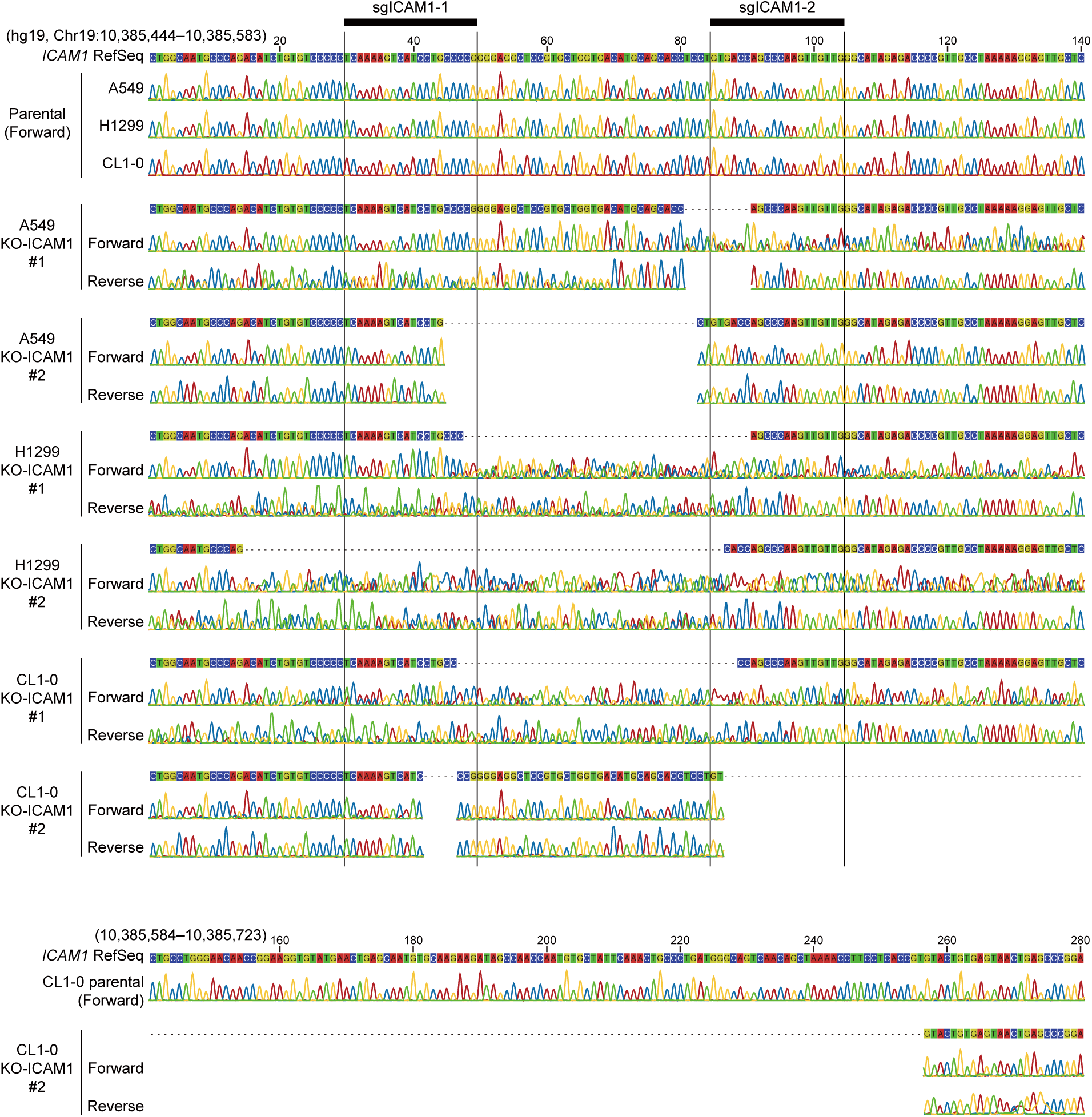
Sanger sequencing validation of the CRISPR/Cas9-edited ICAM1 genome locus of lung cancer cells. Sequencing results of the KO-ICAM1 lung cancer cells are aligned against the reference sequence of the *ICAM1* genome locus. Alignment gaps are denoted as hyphens (-) to mark the lost (knockout) regions of the edited *ICAM1* genome locus.

**Supplementary Figure 8.**
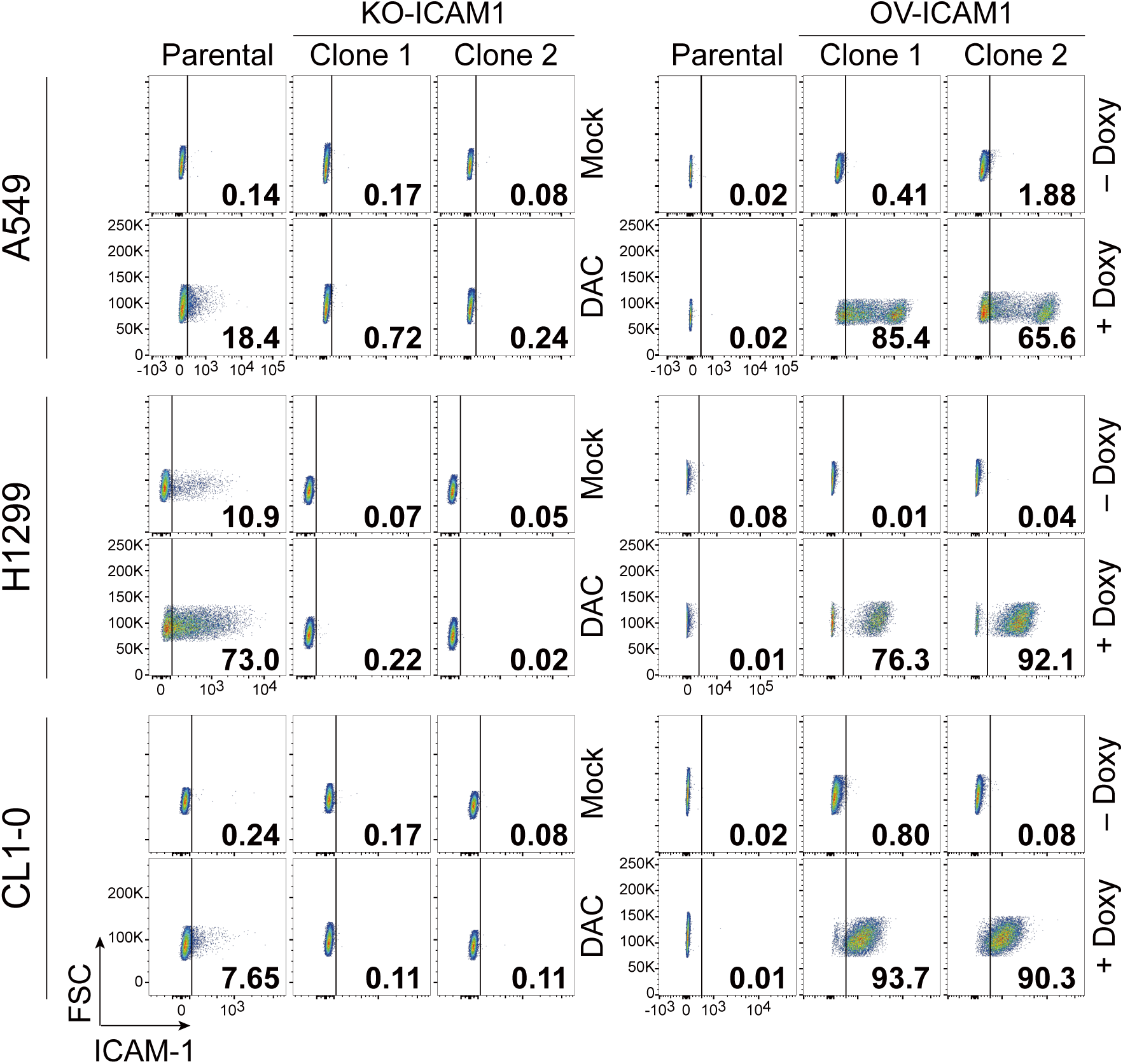
Validation of surface ICAM-1 expression of the ICAM-1-depletion and -overexpression of lung cancer cells. Two independent clones of each genetic manipulation for A549, H1299, and CL1-0 cell lines are shown. X-axis: signal intensities of ICAM-1. Y-axis: forward scatter (FSC).

**Supplementary Figure 9.**
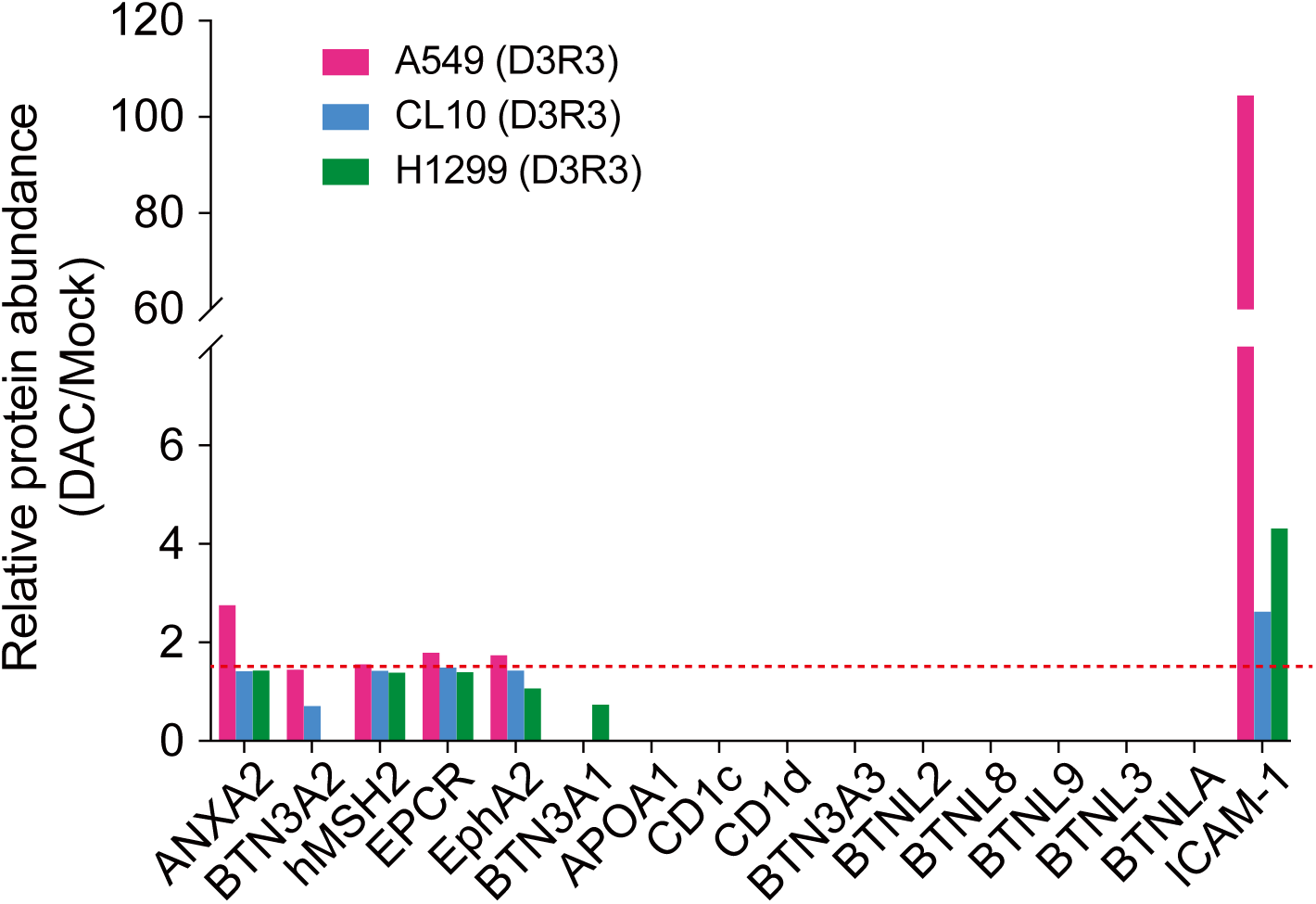
**Relative protein abundance of putative ligands for γδ TCR in surface proteomes of A549, H1299, and CL1-0 cells.**

**Supplementary Figure 10.**
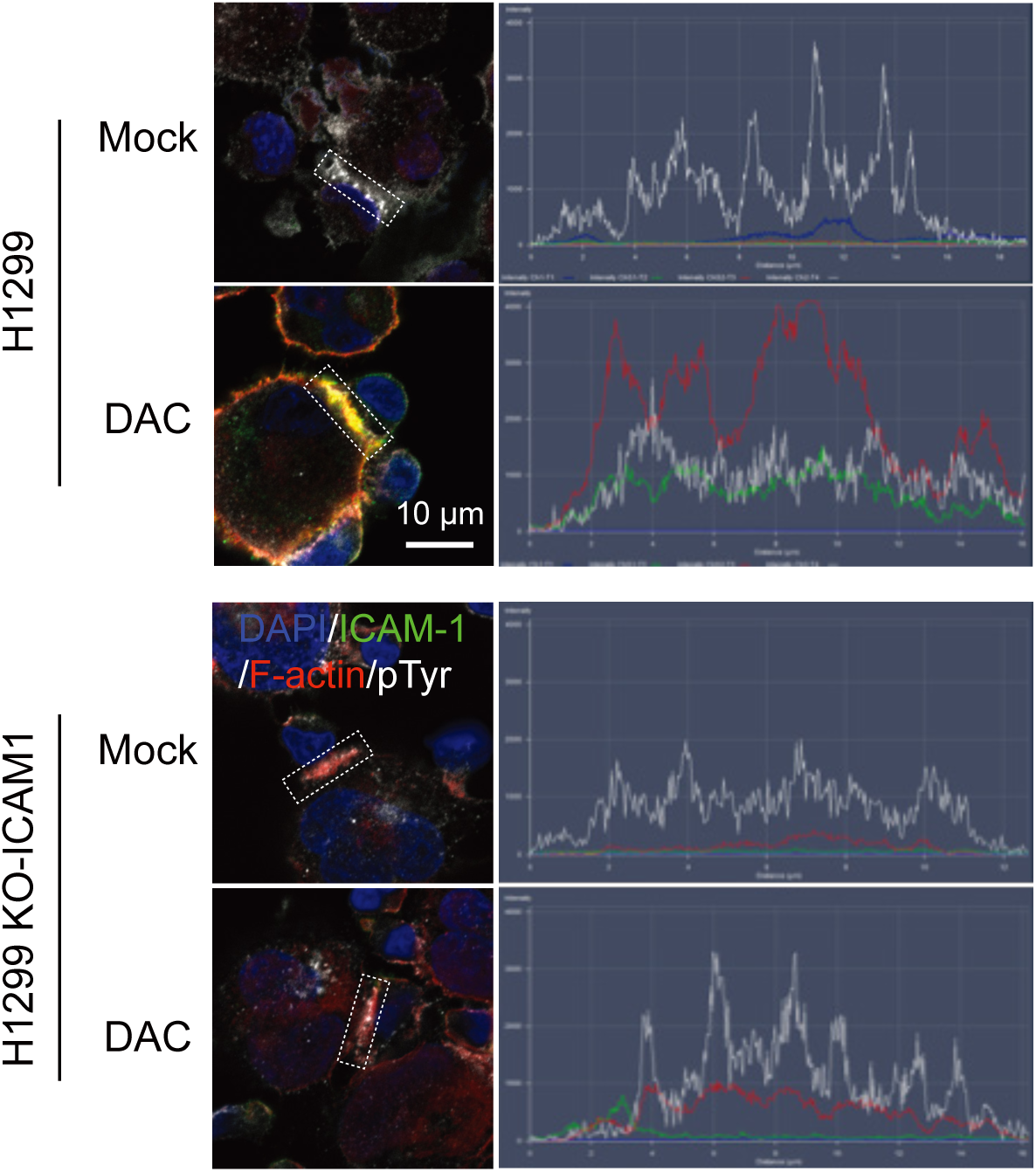
Representative immunofluorescence staining and signal intensity profiles of the proteins at wild-type and ICAM-1 knockout immune synapses between γδ T cells and H1299 lung cancer cells. Parental or ICAM-1 knockout (KO-ICAM1) H1299 cells are pretreated daily with PBS (Mock) or 100 nM DAC for 72 hours followed by 3-day drug-free culture before coculture with γδ T cells. Signal intensities of each protein (F-actin, red; ICAM-1, green; phosphotyrosine, pTyr, white) along the immune synapse area are graphed on the right. DAPI: 4’,6-diamidino-2-phenylindole, as nuclear counterstain. Scale bar: 10 μm.

**Supplementary Figure 11.**
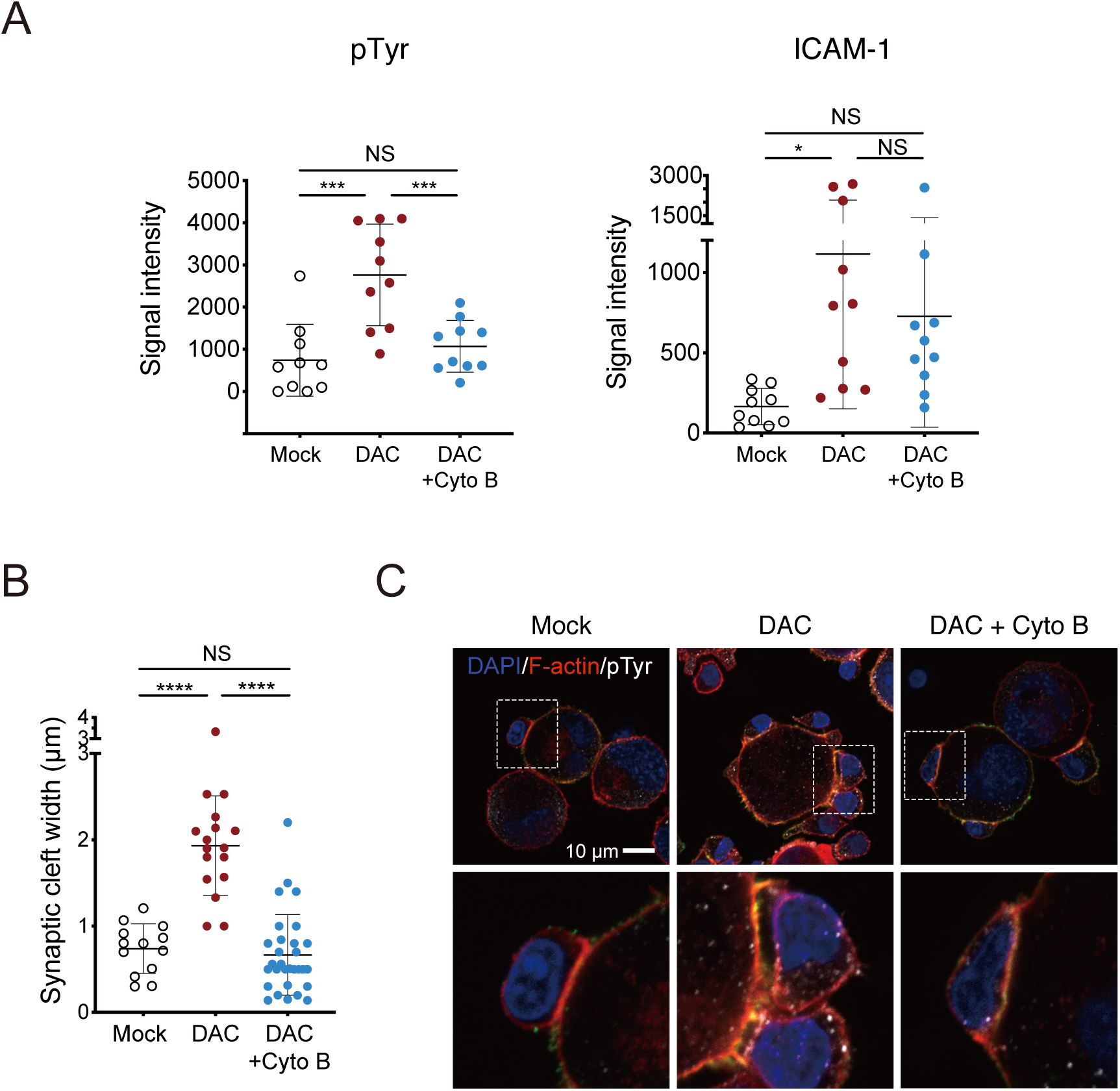
Inhibition of actin polymerization by cytochalasin B disrupts the decitabine-enhanced formation of the active immune synapse. (**A**) Dot plots of pTyr and ICAM-1 signal intensities at immune synapses between γδ T cells and H1299 cells. H1299 cells are pretreated with PBS (Mock), DAC alone or combination of DAC pretreatment (D3R3) and 1 μg/mL cytochalasin B (Cyto B) for 1.5 hours prior to coculture with γδ T cells (mean ± SD). *p* value is calculated by ANOVA with Tukey’s multiple comparisons test (*, *p* < 0.05; ***, *p* < 0.001). (**B**) Dot plots showing the width of immune synaptic cleft between γδ T cells and H1299 cells. H1299 cells are pretreated with PBS (Mock), DAC alone or combination of DAC and Cyto B. Data are presented as mean ± SD. *p* value is calculated by ANOVA with Tukey’s multiple comparisons test (****, *p* < 0.0001). (**C**) Representative immunofluorescence images of synaptic clefts stained for F-actin and pTyr. Scale bar: 10 μm.

**Supplementary Figure 12.**
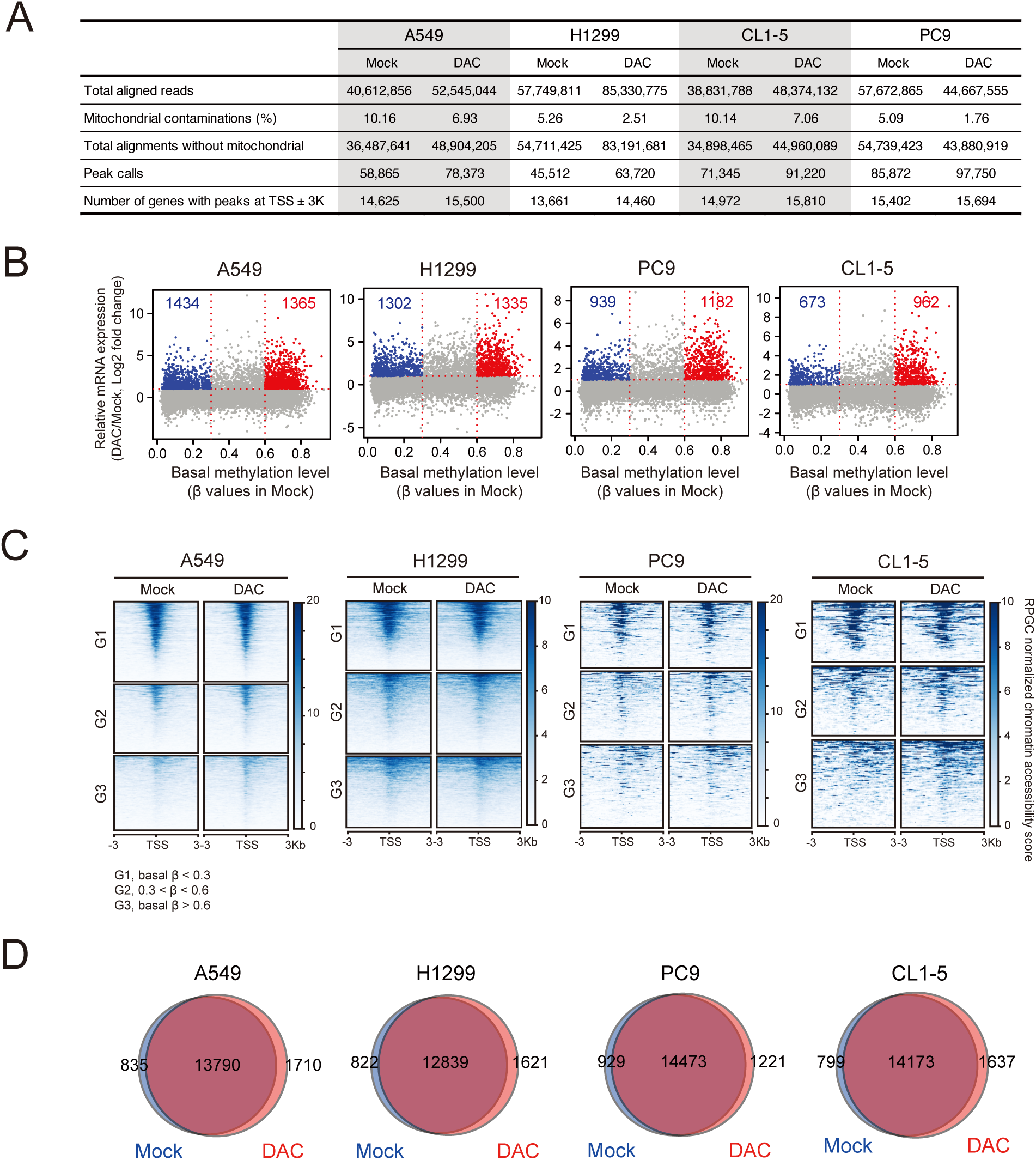
Genome-wide integrative analysis of the fluctuations of genomic DNA methylation and chromatin accessibility status after decitabine treatment. (**A**) The alignment statistics of Omni-ATAC-seq in human lung cancer cell lines. (**B**) Transcriptional changes (y-axis, log2 fold change) in A549, H1299, PC9, CL1-5 cells following DAC treatment at D3R3 and basal methylation levels (x-axis) in cells without DAC treatment (Mock) for all genes measured by mRNA-seq and Infinium MethylationEPIC arrays, respectively. Increases of gene expression by at least 2-fold (Log2 fold change ≥ 1) are considered upregulated by DAC. The methylation levels represent the median β values of promoter probes for each gene. β value = 1, completely methylated; β value = 0, completely unmethylated. (**C**) Promoter chromatin accessibility measured by Omni-ATAC-seq for genes upregulated at least two-fold by DAC in H1299, PC-9, and CL1-5 lung cancer cells. Genes are classified into 3 groups based on β values at baseline (Mock). Chromatin accessibility around transcription start sites (TSS, −3 to +3 kb) are graphed. (**D**) Venn diagram showing the numbers of peaks that represent accessible chromatin at the TSS of all genes in mock-treated vs. DAC-treated lung cancer cells. The broad peak regions of ATAC-seq are called using MACS2 v2.2.5.

**Supplementary Figure 13.**
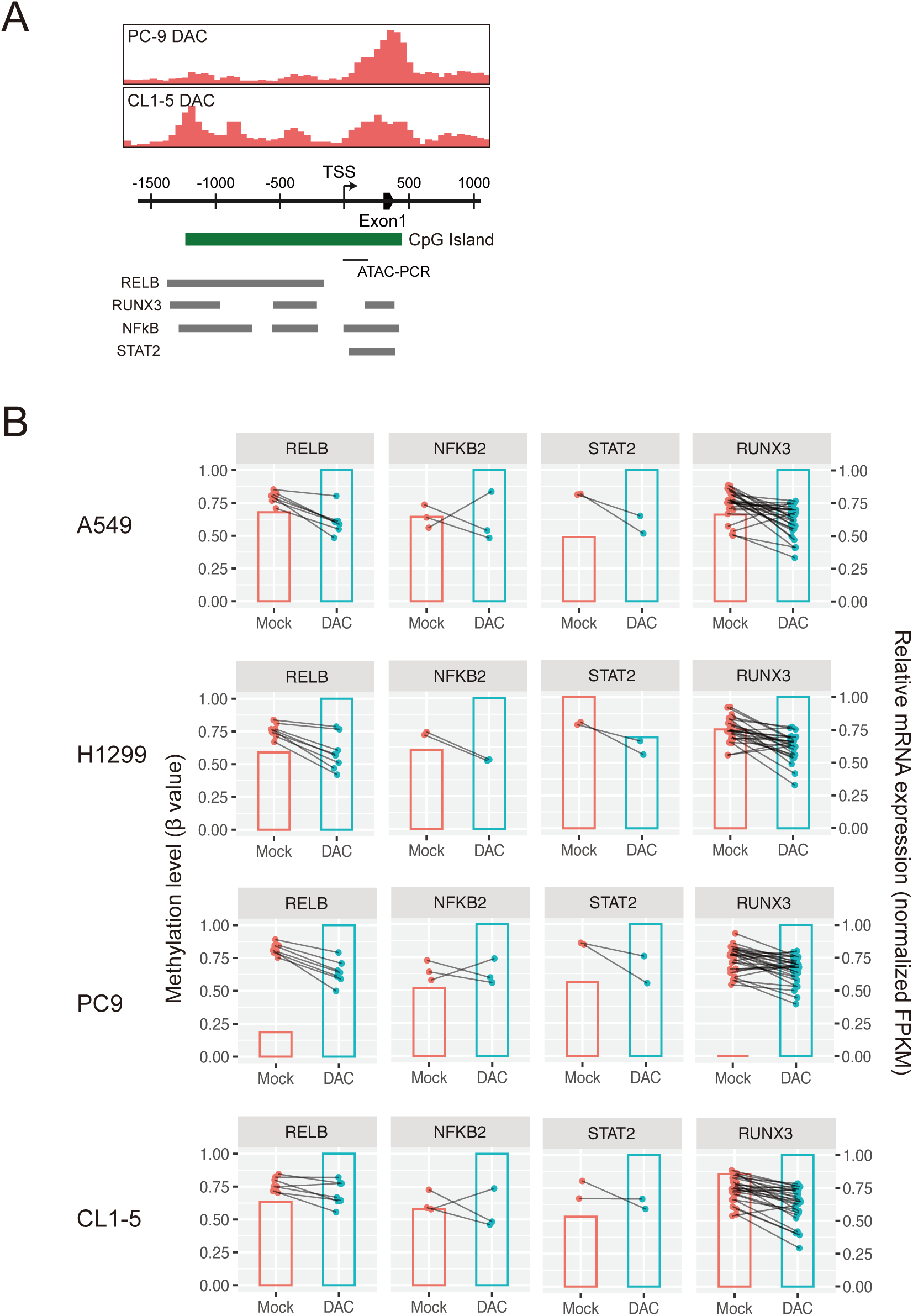
DAC-enhanced ICAM-1 expression may be by via the promoter demethylation of transcription factors targeting the three prime untranslated region (3’-UTR) of *ICAM1* by which upregulate their expressions. (**A**) Diagram of transcription factor binding sites at the *ICAM1* promoter derived from the ENCODE ChIP-seq data (https://www.encodeproject.org). Visualizations of ATAC-seq peaks at the *ICAM1* promoter in PC9 and CL1-5 lung cancer cell lines subject to DAC treatment are shown above. (**B**) Promoter methylation status and mRNA expression levels of putative transcription factors (i.e., RELB, NFKB2, STATS, and RUNX3) at the *ICAM1* promoter in A549, H1299, PC9, and CL1-5 lung cancer cells. Dot and line plots represent methylation levels (β values) of promoter probes measured by Infinium MethylationEPIC arrays. The promoter probes with β values greater or equal to 0.5 at baseline (Mock) are shown. Bar graphs represent relative mRNA expression levels based on normalized FPKM measured by mRNA-seq.

## Notes

### Competing Interest Statement

The authors have declared no competing interest.

